# Mechanisms of Ub-independent ODC recognition and translocation by the 26S proteasome revealing proteasome’s functional adaptability

**DOI:** 10.1101/2025.11.15.688597

**Authors:** Kaijian Chen, Yanxing Wang, Xingyan Ye, Zhanyu Ding, Xuyang Yuan, Cong Xu, Yue Yin, Yao Cong

## Abstract

While most eukaryotic proteins require ubiquitination for proteasomal degradation, a considerable number of substrates undergo ubiquitin-independent degradation. Ornithine decarboxylase (ODC) is a well-known substrate degraded by the 26S proteasome in a ubiquitin-and cofactor-independent manner. However, the underling mechanism of how ODC is recognized and processed without the ubiquitin tag by the 26S proteasome remain unclear. Here, we present eleven cryo-EM structures of the human 26S proteasome in complex with C-terminal truncated or full-length ODC, capturing the entire degradation process from initial recognition to final unfolding. These structures reveal a dynamic, multivalent recognition process involving sequential engagement with vWA domain of Rpn10, Rpt4/5’s coiled-coil, and Rpn2’s PC domain, collectively bypassing the need for ubiquitin. Although the folded ODC core is sufficient for initial binding, its flexible C-terminal tail is crucial for allosterically activating the proteasome’s AAA+ ATPase motor. Furthermore, we also identify a unique translocation gateway where Rpn11’s JAMM motif guides the ODC C-tail into the AAA+ATPase ring, repurposing this deubiquitinase for a novel function. Once activated, the proteasome translocates ODC C-tail and unfolds ODC using its canonical ATP-driven machinery. The continuous structural series from initial substrate recognition through final unfolding elucidates the complete molecular mechanism of ubiquitin-independent ODC degradation.

## Introduction

The 26S proteasome is the principal machinery for selective protein degradation in eukaryotic cells, essential for maintaining cellular proteostasis.^1–5^ Its canonical function relies on recognition of diverse substrates marked by a polyubiquitin chain. These tagged proteins are subsequently unfolded by a powerful AAA+ ATPase motor and translocated into the 20S core particle (CP).^3–5^ Beyond this well-established pathway, the proteasome also degrades a select group of untagged proteins, including many key regulators of growth and proliferation, through ubiquitin-independent pathway.^6–9^ It remains unclear how a machine exquisitely tuned for ubiquitin recognition adapts to process substrates that lack this signal, which is a fundamental question raised by this functional duality. The molecular basis of ubiquitin-independent proteasomal degradation (UbInPD) remains poorly understood.^6,7,9^

The proteasome’s catalytic chamber resides within the 20S CP, a cylindrical barrel-shaped complex, whose access is controlled by the 19S regulatory particle (RP).^3,5,10^ The 19S RP, which is composed of a lid and a base subcomplexes^11^, is responsible for substrate recognition, deubiquitination, and translocation. The base harbors a hexameric AAA+ ATPase motor, a scaffolding subunit Rpn2, and three intrinsic ubiquitin receptors (Rpn1, Rpn10, and Rpn13) that initially capture ubiquitinated substrates.^2,3,5^ Rpn10 recognizes ubiquitin chains via its Ub-interacting motif (UIM) for initial capture.^5,12^ In the lid, the intrinsic deubiquitinase (DUB) Rpn11, part of the JAMM/MPN family DUB, removes the ubiquitin chain to commit the substrate for degradation.^13–20^ The ATPase ring, consisting of N-terminal coiled-coils, OB-ring, and pore-1 loops within the channel, then unfold and thread substrates into the 20S CP by a “hand-over-hand” mechanism powered by ATP hydrolysis.^2,3,5,17,18^

Despite the predominance of the ubiquitin-dependent route, a growing list of physiologically important substrates, such as ODC, p53, c-Fos, FAT10, and Rpn4, are degraded independently of ubiquitin.^21–28^ Recent proteome-wide studies have identified numerous full-length proteins subject to UbInPD.^9^ Some UbInPD substrates require C-terminal degrons and shuttling factors like the Ubiquilin family proteins^9^, while some others are routed through alternative midnolin-proteasome pathway.^8^ UbInPD is thus emerging as an important parallel degradation system, particularly when the ubiquitin availability or conjugation is impaired. However, a mechanistic understanding of how the proteasome directly recognizes and processes non-ubiquitinated substrates remains elusive. Addressing this question is critical not only for understanding proteasomal plasticity but also for advancing ubiquitin-independent targeted degradation strategies with therapeutic targets.

Among the known UbInPD substrates, ornithine decarboxylase (ODC), the rate-limiting enzyme in polyamine biosynthesis^29^, serves as a classical model.^25,26,28^ Polyamines are essential for cell growth and proliferation43. The precise regulation of ODC is crucial, as dysregulated polyamine levels are closely linked to cancer.^30–32^ ODC degradation is mediated by the regulatory protein Antizyme-1 (Az1)^33,34^, which sterically blocks ODC homodimerization and exposes a cryptic proteasome-interacting surface of ODC, thereby facilitating it for ubiquitin-independent degradation by the 26S proteasome.^35^ Notably, a C-terminal tail truncated ODC variant (ODC^ΔC^) can be recognized by the proteasome but not degraded^35^, making the ODC^ΔC^-Az1 complex a valuable model for dissecting substrate recognition in UbInPD.

While cryo-electron microscopy (cryo-EM) has provided high-resolution views of the 26S proteasome engaged with ubiquitinated substrates^12,17,18,36–53^, and those recruited indirectly by cofactors^20,54–61^, a comparable structural understanding of direct, ubiquitin-independent substrate processing has been absent. The key mechanisms, including the intrinsic recognition of folded, non-ubiquitinated proteins, the commitment pathway ensuring affinity and specificity without ubiquitin, and the activation of the ATPase motor, remain to be elucidated.

Here, we address these questions by presenting eleven cryo-EM structures of the human 26S proteasome in complex with ODC-Az1, which together capture the entire process from initial recognition and activation to translocation and final unfolding. Our structures reveal a multivalent, stepwise binding mechanism involving the vWA domain of Rpn10, the Rpt4/5 coiled-coil (CC), and PC domain of Rpn2. We demonstrate that the insertion of ODC’s C-terminal tail into the AAA+ ATPase ring serves as the crucial trigger for proteasome activation, initiating substrate translocation via a conserved “hand-over-hand” mechanism. Strikingly, we discover that Rpn11 uses its JAMM motif to guide initial insertion of the ODC C-terminal tail, revealing an unexpected gateway that allows ubiquitin-independent substrate to bypasses the canonical ubiquitin-binding pathway, and converge on the shared ATP-driven translocation machinery. Together, these findings establish a comprehensive structural and mechanistic framework for UbInPD, expanding our view of the proteasome’s functional plasticity in maintaining proteostasis, and opening new avenues for developing ubiquitin-independent targeted protein degradation for therapeutic targets.

## Results

### Cryo-EM structures of ODC in complex with the 26S proteasome

To structurally characterize how the 26S proteasome recognizes ODC, we employed an established model system comprising C-terminal truncated ODC (ODC^ΔC^) complexed with N-terminal truncated Az1 (Az1^ΔN^) (Fig. S1A-B).^35^ We confirmed that the purified ODC^ΔC^-Az1^ΔN^ complex binds to the human 26S proteasome (Fig. S1D-E), as evidenced by native gel shift assays, showing retarded migration of single- and double-capped proteasomes, western blotting, and mass spectrometry (MS) (Fig. S1F-G, Table S2).

To visualize the recognition process, we determined its cryo-EM structure of the ODC^ΔC^-Az1^ΔN^ bound to the human 26S proteasome after stabilization by glutaraldehyde crosslinking, usually used in macromolecular complex stabilization.^62,63^ This yielded three distinct conformational states of the 26S-ODC^ΔC^-Az1^ΔN^ complex, designated R1, R2, and R3, at 4.4 Å, 4.5 Å, and 3.7 Å resolution, respectively (Fig. 1A, Fig. S2-S3, Tabel S1). The R1 and R2 states were transient, representing 13.7% and 12.7% of the particle population, while R3 state was the predominant (73.6%), more stable conformation (Fig. S2C). As a control, we also determined the structure of the free 26S proteasome at 3.4 Å resolution (Fig. 1A, Fig. S4, Tabel S1), which adopts the characteristic resting conformation.^43,44^ Structural comparison showed that the proteasome in all three 26S-ODC^ΔC^-Az1^ΔN^ maps remained in the resting state. However, a distinct extra density corresponding to ODC^ΔC^-Az1^ΔN^ was observed, located in a canyon formed by the Rpn10, Rpn2, and Rpt4/5 coiled-coil (Fig. 1A-B, Fig. S3D-F). This density became progressively more ordered from the transient R1 state to the stable R3 state (Fig. 1A). We then built an atomic model for each state (Fig. S3G-I). Together, our analysis clearly demonstrates that the proteasome can recognize and bind the ODC core in the absence of its C-terminal tail, but this initial engagement is insufficient to trigger proteasome activation.

**Fig. 1.**
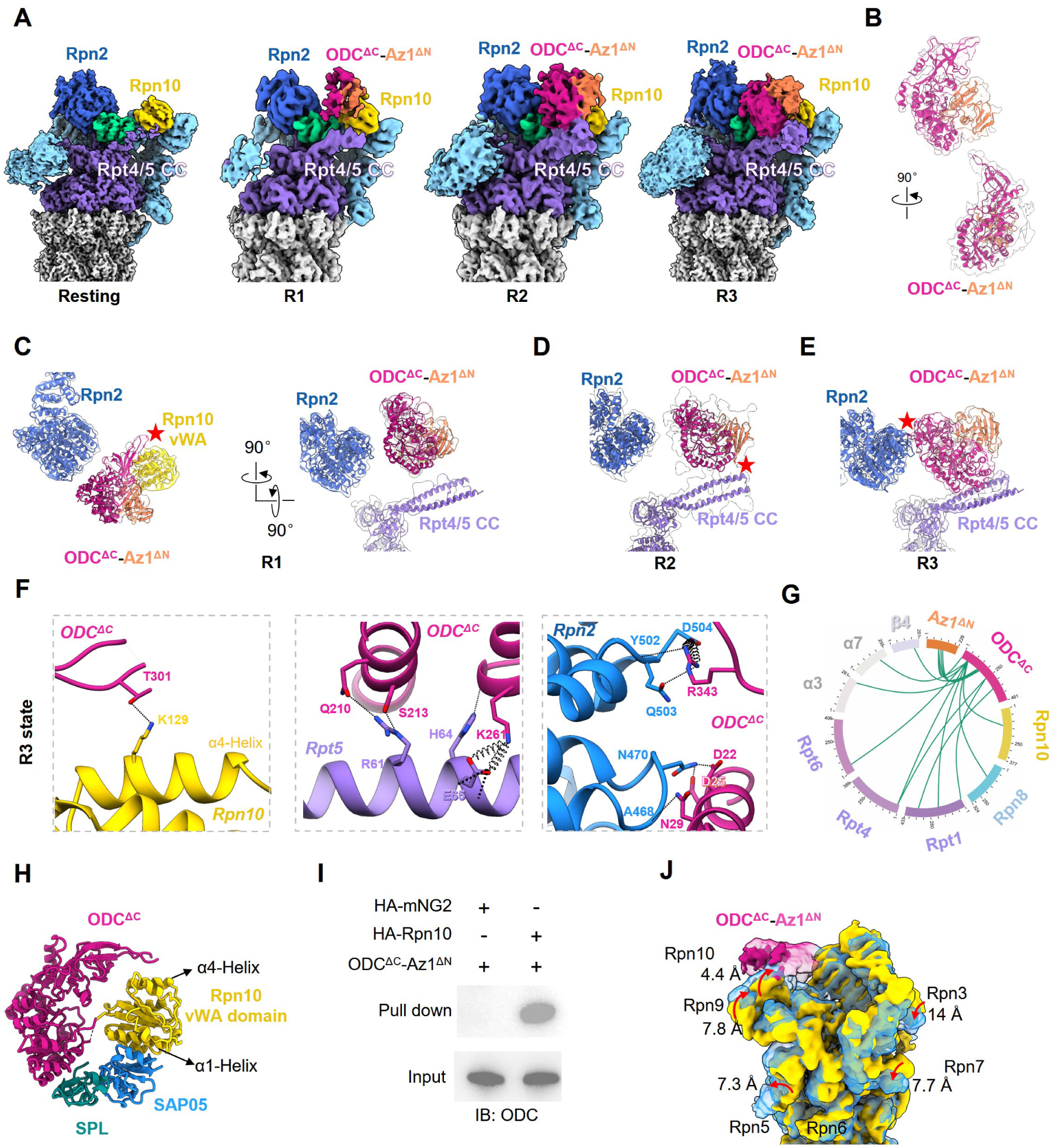
Stepwise recognition of ODC^ΔC^-Az1^ΔN^ by the 26S proteasome. (A) Cryo-EM Maps of the human 26S Proteasome, alone (resting) and in complex with ODC^ΔC^-Az1^ΔN^ corresponding to resting R1, R2 and R3 states. Subunits are colored as follows: lid and Rpn1 (light sky blue), ATPase ring (medium purple), CP (light gray), Rpn10 (gold), Rpn11 (medium spring green), Rpn2 (royal blue), ODC^ΔC^ (medium violet red), and Az1^ΔN^ (coral). This color scheme is followed throughout. (B) Fit of the ODC^ΔC^-Az1^ΔN^ crystal structure (PDB:4ZGY) into the extra density of the R3 state map (transparent surface). (C-E) Sequential engagement of ODC^ΔC^-Az1^ΔN^ with the proteasome. In the R1 state, the substrate density engages only the Rpn10 vWA domain (left, red star), while there is an obvious gap (black arrow) between substrate density and Rpt4/5 coiled-coil or Rpn2 (C). In the R2 state, the substrate engages both Rpn10 and Rpt4/5 coiled coil (D). In the R3 state, the substate is engaged with Rpn10, Rpt4/5 coiled-coil, and Rpn2 (E). (F) Interaction network between ODC^ΔC^ and Rpn10, Rpt5, and Rpn2 in the R3 state. (G) XL-MS analysis of the 26S-ODC^ΔC^-Az1^ΔN^ complex. (H) Structural alignment of the ODC^ΔC^-Rpn10 from R3 state with the SAP05-Rpn10 (PDB: 8PFD) and SPL-SAP05 (PDB: 8PFC). (I) Pull-down assay confirming the interaction between HA-tagged hRpn10 and ODC^ΔC^-Az1^ΔN^. HA-mNG2 is a negative control. Source data are provided as a Source Data file. (J) A representative 3DVA motion of the 26S-ODC^ΔC^-Az1^ΔN^ dataset, showing the two extreme conditions in the variance, with directions of the subunit motion indicated.

### A stepwise, multivalent binding mechanism for ODC recognition

Our structure analysis of the R1 to R3 states, supported by 3D variability analysis (3DVA), reveals a dynamic, stepwise mechanism for ubiquitin-independent substrate recognition (Fig. 1C-E, J, Movie 1). The process begins in the R1 state, where the ODC β-sheet domain makes its initial contact with the vWA domain of Rpn10 and is resolved (Fig. 1C, Fig. S3J-K). Then the complex transitions to the R2 state, where the ODC α/β barrel domain becomes better resolved (Fig. S3E) and docks onto the Rpt4/5 coiled-coil, primarily engaging Rpt5 (Fig. 1D). Rpt5 has been reported as a receptor for polyubiquitin chain20, and recent structural studies confirmed that Rpt4/5 coiled-coil directly interacts with K48-linked ubiquitin, the primary signal for proteasomal degradation16. This engagement induces an ordering of the otherwise intrinsically flexible Rpt4/5 coiled-coil (Fig. 1A). The recognition sequence culminates in the stable and better resolved R3 state, with the crystal structure of ODC^ΔC^-Az1^ΔN^ fitting well into the substrate density^35^ (Fig. 1B), where the ODC-Az1 complex is full engaged, now also locked into place by interactions with the PC domain of Rpn2 (Fig. 1E).

Structural analysis identified intricate network of salt bridges and hydrogen bonds (H-bonds) that stabilizes the ternary interface, further corroborated by our chemical cross-linking mass spectrometry (XL-MS) data (Fig. 1F-G). It appears that the initial ODC binding is mediated by the α4-helix of the Rpn10 vWA domain (Fig. 1H). In contrast, a recent study suggests that Rpn10 recognizes the plant parasite cofactor SAP05 via Rpn10 α1-helix, enabling indirect capture of Ub-independent substrates SPL and GATA developmental regulators in Arabidopsis thaliana (Fig. 1H).^64–66^ Our findings, along with recent studies^64–67^, suggest that Rpn10 serves as an important Ub-independent substrate receptor via its vWA domain, with distinct recognition elements for different substrates/cofactors. Furthermore, our in vitro pull-down assays confirmed the Rpn10-ODC interaction (Fig. 1I).

Moreover, our 3DVA visualizes the entire process as a continuous conformational rearrangement, where the initial ODC binding event at Rpn10 and the Rpt4/5 coiled-coil triggers a considerable clockwise rotation of the 19S lid (Fig. 1J, Movie 1), dynamically orienting the substrate toward and eventually engagement with the Rpn2 PC domain. This reveals how proteasome repurposes its canonical ubiquitin-recognition machinery for direct, high-affinity capture of an untagged substrate. Remarkably, despite these large conformational changes in the 19S lid and the Rpt4/5 and Rpt3/6 coiled-coils, the proteasome remains in an inactive, resting state, characterized by a closed 20S gate (Fig. S3L).

### Multivalent binding mechanism of ODC to proteasome and the essential role of its C-terminal tail in proteasome activation

While the ODC core is sufficient for recognition by proteasome, it is not degraded. To determine the role of the ODC C-terminal tail in proteasome activation and subsequent substrate translocation, we prepared a complex of full-length ODC and Az1 (Fig. S1A, C). It can readily be degraded by proteasome in vitro (Fig. S1H), confirming the functional activity of our model system, consistent with previous works.^26,33^ We then performed cryo-EM analysis of the full-length ODC-Az1 in complex with 26S proteasome without cross-linking. In sharp contrasts to the ODC^ΔC^ sample, which produced all resting state (Fig. 1A), the full-length ODC sample drove approximately half of the proteasome population into an activated conformation, while the other half remained in resting state (Fig. S5). This provides direct evidence that ODC’s flexible C-terminal tail is the essential trigger for proteasome activation and subsequent substrate processing.

Through multiple rounds of 3DVA and 3D classification, we resolved eight distinct cryo-EM maps of the 26S-ODC-Az1 complex: three resting states (B1-B3, at 3.4 Å, 3.6 Å, 3.9 Å resolution; Fig. 2A, Fig. S5, and Table S1), and five activated states (A1-A5, 3.4 to 4.4 Å resolution; Fig. 3A, Fig. S6, and Table S1). We built an atomic model for each map, which matched well with the corresponding density map (Fig. S7). In the resting B1-B3 states, the presence of the C-terminal tail appeared to stabilize their interaction, resulting in better-resolved ODC-Az1 compared to that of the tail truncated substrate. The overall B1 conformation is comparable to the R3 state with minor confirmational changes in the lid (Fig. S8A), whereas the transient R1 and R2 states were not detected.

**Fig. 2.**
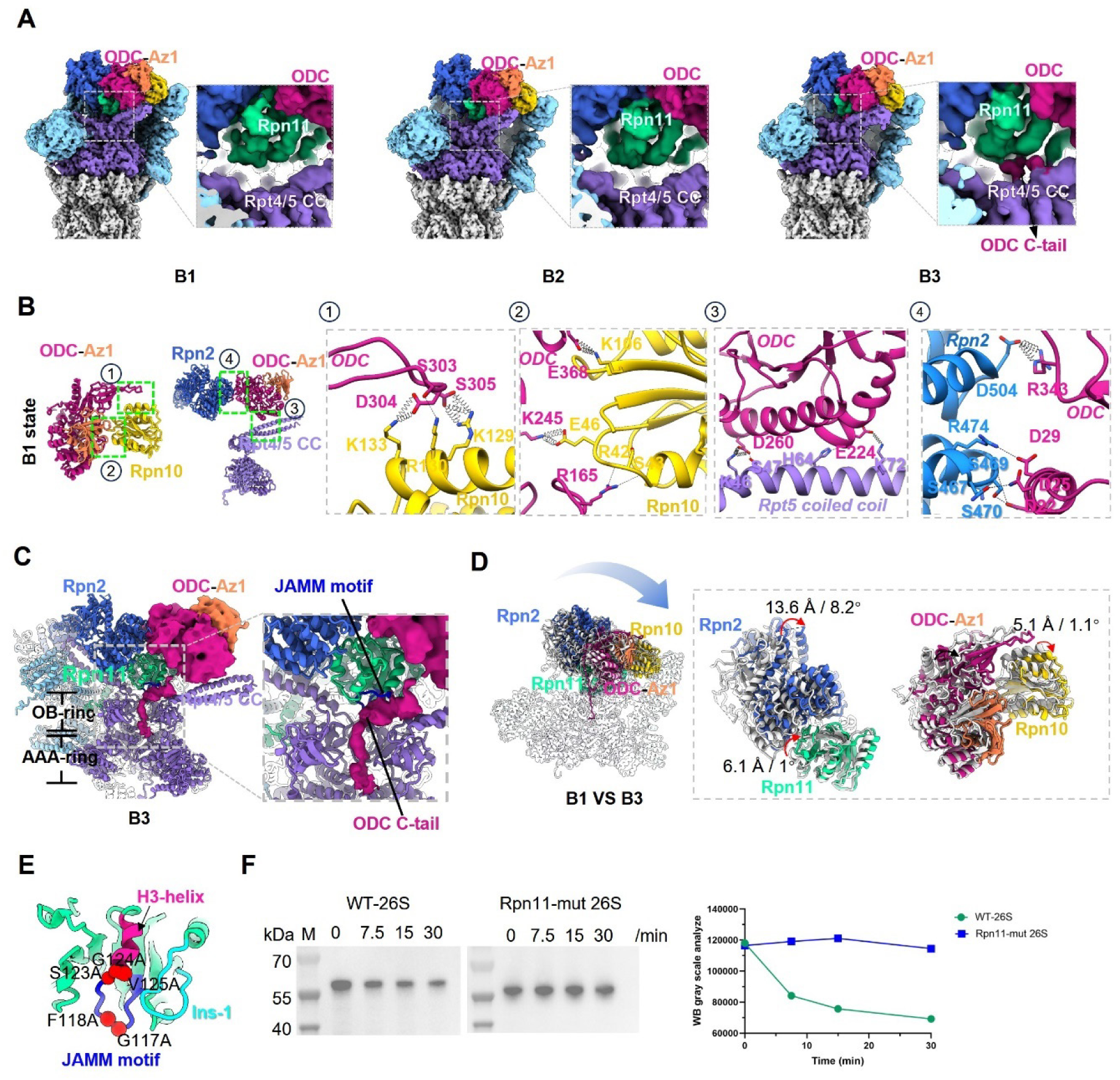
The initial insertion of ODC C-terminal tail into the ATPase ring. (A) Cryo-EM maps of the 26S proteasome in complex with full-length ODC-Az1, showing the B1, B2, and B3 resting states. A zoomed-in view highlights a new density for the ODC C-terminal tail (medium violet red) appearing between Rpn11 and the Rpt4/5 OB-ring in the B3 state. (B) Interaction network between full-length ODC and Rpn10/Rpn2/Rpt5 in the B1 state. (C) The ODC C-terminal tail (density in medium violet red) interacts with the Rpn11 JAMM motif (in blue) and the Rpt4/5 OB-ring gate in the B3 state (density from unsharpened map). ODC C-tail density was colored in medium violet red. (D) Superposition of the B1 (gray) and B3 (colored) states, revealing conformational variations in key subunits. (E) Location of mutated residues (red spheres) in the Rpn11 JAMM motif. The other key structural elements are also indicated: H3-helix in pink, and Ins-1 in cyan. (F) Western blot analysis of an *in vitro* degradation assay. Proteasomes containing the Rpn11 JAMM mutant fail to degrade ODC. Source data are provided as a Source Data file.

**Fig. 3.**
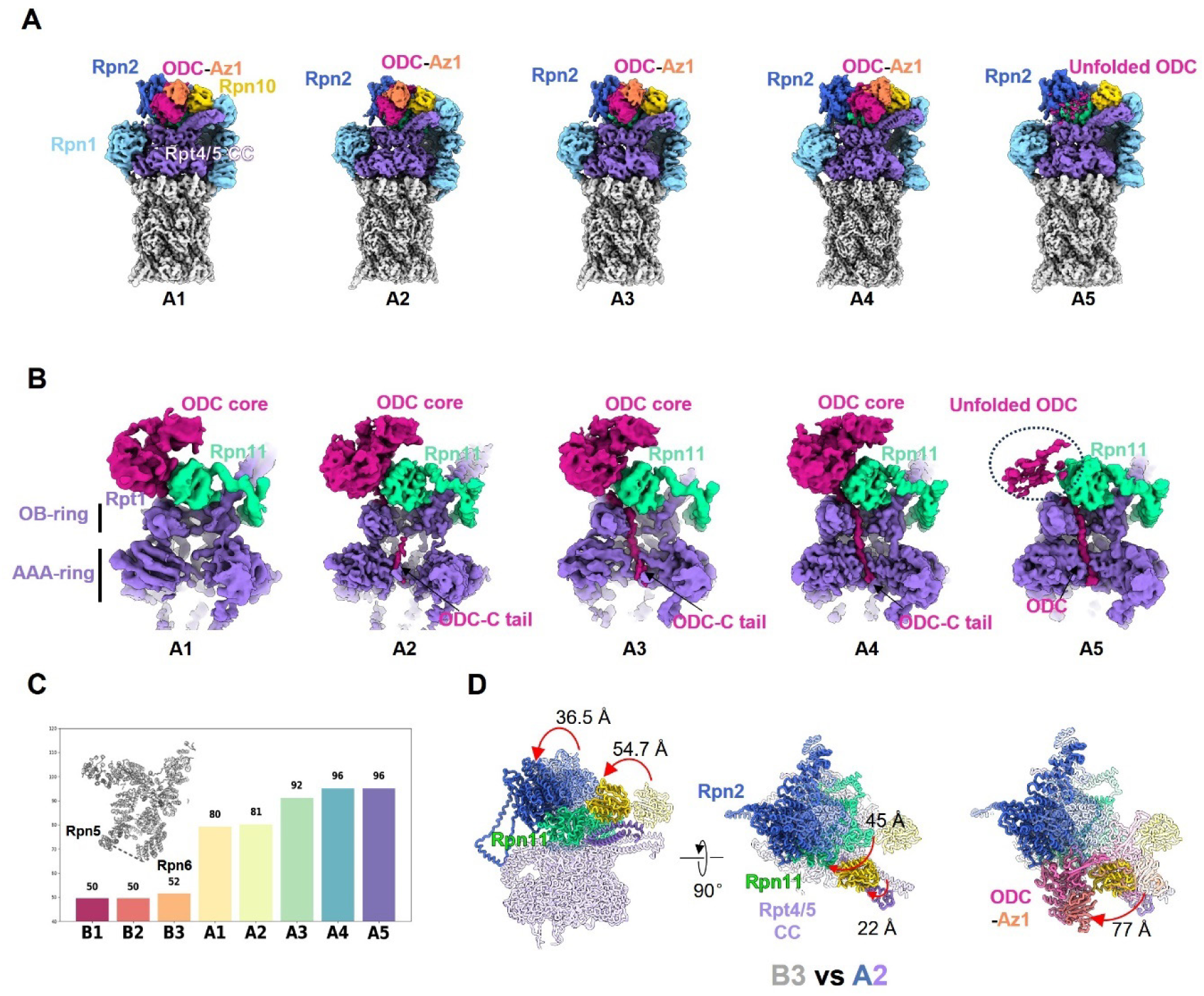
ODC C-terminal tail insertion activates the 26S proteasome. (A) Cryo-EM maps of the full-length ODC-Az1 bound 26S proteasome in five distinct activated states (A1-A5). (B) Cutaway views of ATPase-ring showing the ODC C-terminal tail progressive threaded through the central channel from state A1 to A5. The density for Rpt4/5 has been omitted for clarity. (C) The distance between Rpn5 and Rpn6 increases across the resting (B1-B3) and activated (A1-A5) states, indicating a continuum of activation. (D) Superposition showing the large-scale translation of ODC-Az1 and associated proteasome subunits (Rpn2, Rpn10, Rpn11, Rpt4/5) during the transition between B3 and A2 state.

Detailed analysis of the better-resolved B1 state structure reveals the full-length ODC maintains multivalent contacts with the 26S proteasome. The ODC-Rpn10 interaction is slightly enhanced in the B1 state compared to the R3 state: in addition to the U-loop (residues E299-E310) of ODC contacting the α4-helix of Rpn10 vWA domain (Fig. 1F, 2B), an additional contact formed between the ODC α/β barrel domain and the α1-helix of the Rpn10 (Fig. 2B), a structural element also involved in the SAP05 recognition^64–66^ (Fig. 1H). Moreover, the α7/8-helix of the ODC α/β barrel domain engages the Rpt4/5 coiled-coil (specifically, Rpt5’s K46, S47, H64, and K72), and its L340-E348 loop forms extensive networks of H-bonds and salt-bridges with the T1-/T2-like sites of the Rpn2 PC domain (Fig. 2B, S3J). These extensive, multivalent interactions, which are maintained across the B1-B3 states, demonstrate how the proteasome has evolved to achieve high-affinity binding of a UbInPD substrate without a ubiquitin signal, securely anchoring the substrate in the repurposed Rpn10-Rpn5-Rpt2 canon, in preparation for its activation and subsequent processing.

### Rpn11 guides ODC C-terminal tail into the AAA+ motor through its JAMM motif

Comparison of the B1-B3 resting states reveals the trajectory of the ODC C-terminal tail as it prepares for insertion into the AAA+ motor. In the B3 state, an additional density corresponding to the ODC C-terminal tail emerges in a tunnel between Rpn11 and the Rpt4/5 coiled-coil, extending toward the OB-ring gate (Fig. 2C). This “initial binder” (encompassing up to the middle C-tail region) segment of the tail interacts with the JAMM motif of Rpn11, the base of the Rpt4/5 coiled-coil, and the OB-ring (Fig. 2C). The B3 state thus captures the initial insertion of the C-terminal tail, a critical step for committing ODC to degradation. This insertion is accompanied by coordinated conformational changes in the adjacent Rpn2, Rpn10, and Rpn11 across the B1 to B3 states (Fig. 2D). Specifically, Rpn2 rotates clockwise up to 13.6 Å, and Rpn10 rotates in the same direction by 5.1 Å. These motions drive a 6.1 Å upward shift of Rpn11, positioning its JAMM motif to capture the incoming ODC tail (Fig. 2C-D).

Notably, we find that for ODC, its initial binder engages the JAMM motif of Rpn11 (Fig. 2C), in sharp contrast to Ub-dependent substrates, where ubiquitin chain engages the Insert-1 β-hairpin of Rpn11 for deubiquitination (Fig. 4B).^14,15,17,18^ To validate the functional importance of this newly defined interaction, we mutated key interacting residues in the JAMM motif (G117, F118, S123, G124, and V125) to alanine (Fig. 2E). Degradation assays revealed that proteasomes containing this mutant Rpn11 was unable to degrade ODC-Az1 (Fig. 2F), demonstrating the essential role of the Rpn11 JAMM motif in stabilizing the initial ODC C-tail engagement. This finding highlights how a UbInPD substrate has evolved to exploit the proteasome’s architecture, repurposing the canonical deubiquitinase Rpn11 to provide a new gateway to guide its C-terminal tail into the ATPase motor for degradation.

**Fig. 4.**
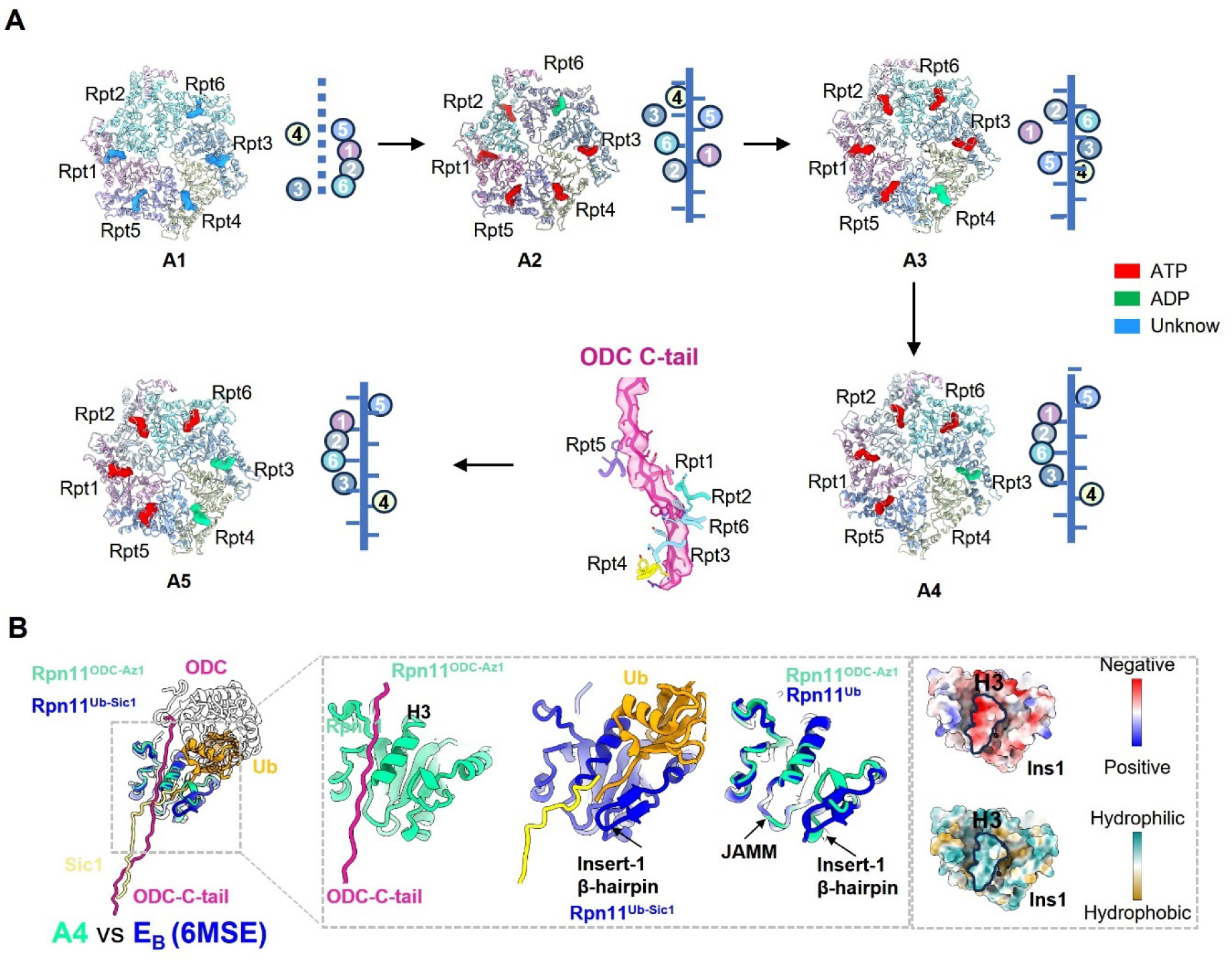
ODC translocation is driven by the conserved AAA+ ATPase motor and a distinct Rpn11 gateway for UbIdPD substrate ODC. (A) Top views of the ATPase ring (ribbon representation), with the nucleotide state (ATP, red; ADP, green; unknown, blue) of each Rpt subunit indicated. Showing also the spiral staircase of pore-1 loops engaging the ODC C-terminal tail during translocation (A1-A4) and unfolding (A5). The A1 state is less well resolved and the nucleotide state is uncertain to assign (colored in light blue). A detailed view of the A4 state shows the ODC tail (density and model in medium violet red) securely gripped by the pore-1 loops of all six Rpt subunits, with an additional contact from the Rpt5 pore-2 loop. (B) Comparison of Rpn11 engagement in ubiquitin-independent (A4 state) and ubiquitin-dependent (E_B_ state, PDB: 6MSE) degradation. In the A4 state (left in zoomed-in view), the ODC C-terminal tail (medium violet red) interacts with the H3-helix rim of Rpn11 (medium spring green). This is contrasted with a canonical substrate-engaged state (middle; PDB 6MSE), where ubiquitin (orange) binds to the Insert-1 of Rpn11 (blue). The structural alignment (right) of Rpn11 reveals that in the ODC-bound state, the Insert-1 β-hairpin undergoes a conformational switch, moving inward to lock the canonical ubiquitin gateway. Corresponding electrostatic surface views are also shown.

### ODC C-terminal tail insertion triggers proteasome activation and coordinated substrate translocation

The insertion of the ODC C-terminal tail into the motor unleashes a cascade of conformational changes that transform the proteasome into a fully active, degradation-competent state. Across the five activated states we resolved (A1-A5) (Fig. 3A), we observe the canonical hallmarks of activation: the C-terminal HbYX motifs of Rpt1/2/3/5/6 insert into the α-ring pockets to open the 20S gate (Fig. S9B), while the tail insertion into the ATPase ring induces a significant clockwise rotation of the lid and downward movement of the OB-ring, aligning the ATPase channel with the 20S CP entrance to form a contiguous path for the substrate (Fig. 3B, Fig. S8B). These states represent a continuum of activation, characterized by a progressive increase in the distance between Rpn5 and Rpn6 (from ∼80 Å in A1/A2 states to 96 Å in A4/A5 states) (Fig. 3C, Fig. S9A), which reflects the dynamic motions that facilitate substrate unfolding and translocation, comparable to what has been observed for Ub-dependent substrates^17,18^. Critically, while the folded ODC core remains bound and ordered across the A1-A4 states, its C-terminal tail, initially disordered in state A1, becomes progressively better defined in A2-A4 states, signifying its progressive engagement and movement through the AAA+ motor (Fig. 3B).

A1 and A2 share similar architectures and Rpn5-Rpn6 spacing, but the A2 map is better resolved, exhibiting a visible ODC tail (Fig. 3A-B, Fig. S8D). A comparative analysis reveals a transition from the pre-activation B3 state to the activated A2 state involves a pronounced rotation of the 19S RP (Fig. 3D). The lid undergoes significant clockwise rotation, in which process, Rpt4/5 coiled-coil rotates less (22 Å) than Rpn2 and Rpn10 (36.5 Å / 54.7 Å), releasing ODC from Rpt4/5 while it moves together with Rpn2 and Rpn10 (Fig. 3D). In a concerted motion, the ODC-Az1 rotates in the same direction by up to 77 Å, moving along with Rpn10 and the underlying Rpn11 (Fig. 3D). This movement disrupts the contact between ODC and the Rpt4/5 coiled-coil while strengthening its association with Rpn2. This hand-off effectively transfers the substrate from its initial recognition cradle and positions its core in immediate proximity to the AAA+ motor, with its C-terminal tail now threaded into the translocation channel for subsequent processing (Fig. 3B). Further progression from A2 to A4 states involves a 17 Å translation of ODC-Az1 in the opposite direction, while Rpn2 and Rpn11 undergo only minor movements (Fig. S8E), representing a final structural adjustment that facilitates efficient substrate processing.

### ODC degradation converges on the conserved hand-over-hand translocation mechanism

Once the ODC C-terminal tail is inserted and the ATPase-ring activated, the AAA+ motor progressively threads the ODC tail through its central channel using the conserved, ATP-hydrolysis-driven “hand-over-hand” mechanism, previously observed in ubiquitinated substrates (Fig. 4A).^5,17,18^ Our structures capture this entire process, beginning with the transition from the B3 resting state to the A1 activated state, where the descent of the Rpt3 pore-1 loop from the highest to the lowest position pulls the ODC tail into the motor, triggering its activation (Fig. 4A, Fig. S10A). In the A1 state, once deeper insertion of the tail allows it to touch the Rpt3/4 pore-1 loops (Fig. S8F), this triggers ATP hydrolysis and allosterically activated proteasome complex. The A1 state represents a transient, early activation intermediate poised for translocation, characterized by dynamic interactions and poorly resolved nucleotide densities.

From this initiation state, the proteasome progresses through A2 to A4 states, coupling sequential ATP hydrolysis events around the Rpt ring to the progressive translocation of the ODC tail deeper into the ATPase channel (Fig. 3B, 4A). During A2 to A3 transition, Rpt6 nucleotides exchange from ADP to ATP, while Rpt4 hydrolyzed ATP to ADP. The progression from A3 to A4 transition involves Rpt3 hydrolyzed ATP to ADP and ADP release in Rpt4 (Fig. 4A, Fig. S10B). This translocation culminates in the A4 state, which closely resembles the canonical E_D2_ state, in the proteasome processing of the K63-ubiquitinated Sic1 substrate^18^ (Fig. S8G). However, a key distinction emerges: whereas some ubiquitinated substrate are engaged by five-loop (except Rpt5) as in the E_D2_ state^18^, the ODC tail is gripped by all six pore-1 loops in our A4 state (Fig. 4A). This suggests a more secure grip on ODC, priming it for unfolding. This pre-unfolding state is followed by the A5 state, where the globular density of the ODC core becomes diffuse (Fig. 3A-B), and a continuous density of the unfolded polypeptide is visible extending throughout the ATPase channel, likely representing an unfolded state (Fig. 3B). This continuum of structures (A1-A5) provides an unprecedented, high-resolution timeline of Ub-independent degradation, from initial molecular touch that awakens the proteasome to final substrate unfolding.

Our structures also reveal that Rpn11 is repurposed not only for initial tail insertion but also as a novel anchor point during translocation. In the canonical pathway, the Insert-1 (Ins1) β-hairpin of Rpn11 engages the ubiquitin chain via a hydrophobic pocket (Fig. 4B), positioning it for cleavage.^17,18^ Our well-resolved A4 state shows the ODC C-terminal tail completely bypassing this site. Instead, the tail forms a distinct interaction with the rim of the H3-helix of Rpn11 (Fig. 4B). This previously unidentified contact point serves as a non-canonical anchor, stabilizing the ODC tail during translocation, in the absence of a ubiquitin handle. This finding underscores a remarkable plasticity of the proteasome, which has repurposed its DUB Rpn11 to engage an untagged substrate through two distinct, novel interfaces, one for insertion and another for translocation. Such mechanistic divergence ensures efficient processing of diverse clients to maintain proteostasis.

## Discussion

The ubiquitin-proteasome system is central to eukaryotic protein homeostasis, yet the molecular principles that govern the degradation of substrates that bypass the canonical ubiquitination pathway have remained obscure. ODC, a key enzyme in polyamine biosynthesis, serves as a classical model for UbInPD, providing a unique opportunity to address this knowledge gap. In this study, we present eleven distinct cryo-EM structures of the human 26S proteasome in complex with ODC-Az1, capturing the entire process from initial substrate recognition and proteasome activation to translocation and final unfolding. Our work reveals a sophisticated mechanism centered on three core features: a multivalent recognition and staged activation mechanism that compensates for the absence of ubiquitin (Fig. 1-3, Movie 1); the repurposing of the deubiquitinase Rpn11 as a novel gateway for substrate commitment (Fig. 2C, E-F, 4B); and the ultimate convergence of this unique pathway on the proteasome’s conserved ATP-driven translocation machinery (Fig. 4A). These findings illuminate the molecular basis of UbInPD, the proteasome’s remarkable functional plasticity, and open new avenues for the development of targeted protein degradation strategies.

### A new pattern for substrate recognition: multivalency bypasses ubiquitination

A central question in UbInPD is how the proteasome achieves specificity and affinity for a substrate without a ubiquitin tag. Our structural analysis reveal that ODC degradation is initiated by a stepwise, multivalent recognition strategy that leverages allostery to generate high-affinity recognition to compensate for the absence of ubiquitin tag (Fig. 5). The process initiates with the ODC core binding to the vWA domain of Rpn10, which serves as the primary landing pad (Fig. 1A, C-E, Movie 1). This event triggers a cascade of conformational changes, inducing the ordering of the flexible Rpt4/5 coiled-coil to create a secondary binding site (specifically with Rpt5). These allosteric movements propagate through the ATPase OB ring to the Rpt3/6 coiled-coil, where Rpn2 is anchored, inducing clockwise lid rotation that orients ODC toward Rpn2 PC domain. This final engagement locks the substrate into a fully engaged, pre-degradation state. Eventually, in the B3 state, the ODC C-terminal tail contacts the Rpn11 JAMM motif (Fig. 2C), inserting into the OB-ring but not yet reaching pore-1 loops. This commits ODC to degradation without ubiquitin, bypassing traditional deubiquitination. This extensive interface, which repurposes at least three canonical ubiquitin-recognition modules (Rpn10, Rpn2, and Rpt5), demonstrates how cooperative, multivalent engagement can generate the affinity and specificity required to bypass the need for a ubiquitin signal.

**Fig. 5.**
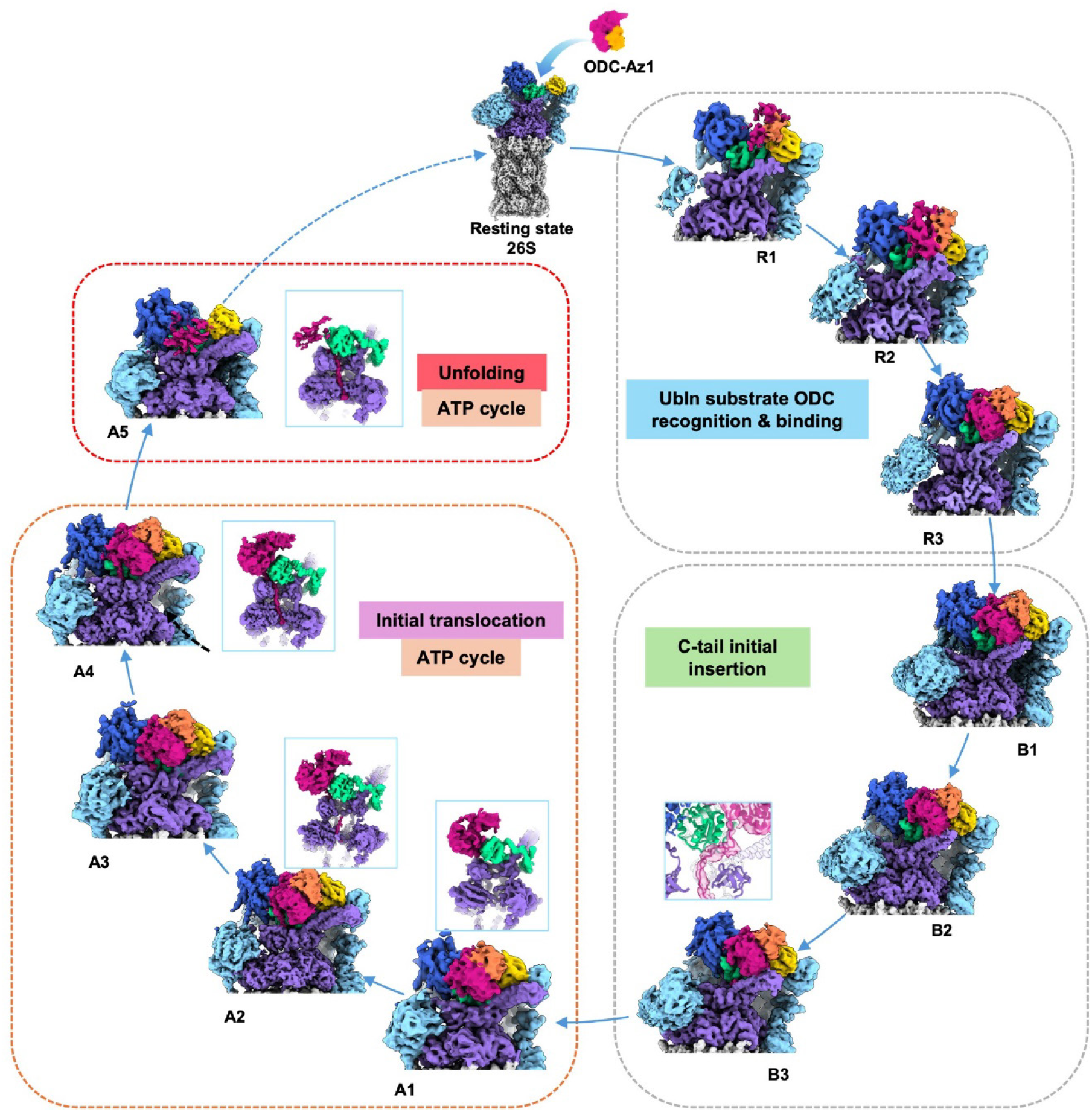
Proposed the mechanism for ubiquitin-independent recognition and degradation of ODC by the 26S proteasome. The process begins with the stepwise, multivalent recognition of the ODC core, starting with Rpn10 (R1), followed by Rpt4/5 coiled-coil (R2), and finally Rpn2 (R3). In the presence of the ODC C-terminal tail (B1-B3 states), conformational changes in the lid eventually position the Rpn11 JAMM motif to capture the tail (B3). Insertion of the ODC tail into the ATPase motor triggers proteasome activation (A1), followed by translocation of the polypeptide via ATP hydrolysis (A2-A4) and, ultimately, unfolding of the ODC core (A5).

### The C-terminal tail triggers proteasome activation and coordinated translocation

ODC core binding maintains proteasome in a resting state (Fig. 1A), but the C-terminal tail is essential for activation, driving approximately half of proteasomes into activated conformations (Fig. S5C). The trigger for this activation appears to be the insertion of the tail deep enough into the motor to touch the Rpt3/4 pore-1 loops in the A1 state. This triggers ATP hydrolysis and allosterically activates the entire complex (Fig. 3, 5). Consistent with this model, a recent study has shown that inhibiting the Rpt3/4 pore-1 loop impairs 19S activation.^57^ Thus, the ODC C-terminal tail functions as a bipartite signal that mediates both substrate commitment and proteasome activation.

Once activated, the ODC tail is griped by the pore-1 loops of the motor, enabling its progressive translocation (Fig. 4A, 5). During the initial transition to the A2 state, a significant clockwise rotation of the lid repositions the substrate, releasing it from the Rpt4/5 coiled-coil while enhancing its interaction with the Rpn2 PC domain (Fig. 3D). As translocation proceeds to the A4 state, the ODC tail’s contact with Rpn11 shifts from the JAMM motif to the H3-helix rim, and all six pore-1 loops firmly grip the ODC tail preparing for unfolding (Fig. 2C, 4). This six-loop engagement suggests a more secure grip compared to the five-loop engagement observed for some ubiquitinated substrates.^17,18^ Finally, in the A5 state, the diffuse density of the ODC core indicates unfolding intermediates, with continuous unfolded polypeptide density extending through the ATPase ring, likely representing an unfolded state (Fig. 3B). These observations show that despite distinct recognition and entry mechanisms, both pathways converge on the proteasome’s canonical ATP-driven translocation machinery, employing a conserved “hand-over-hand” mechanism, highlighting the proteasome’s adaptability to diverse substrates.

### The repurposed deubiquitinase Rpn11 provides a new gateway for UbInPD degradation

Our study reveals that Ub-independent ODC degradation diverges from the canonical pathway at the initial substrate recognition sites and commitment steps. The canonical pathway relies on a network of ubiquitin receptors, including the Rpn10 UIM motif, the Rpt4/5 coiled-coil (for K48-linked chains), and Rpn2 (for K11-linked chains).^12,53^ In contrast, ODC engages the proteasome directly via a multivalent strategy that repurposes these elements. ODC utilizes:

1. the Rpn10 vWA domain (α4- and α1-helices) for direct binding to its core; (2) the Rpt4/5 coiled-coil (specifically Rpt5) as a transient anchor that is released upon activation; and (3) the Rpn2 PC domain (T1/T2-like sites) to lock the substrate in place. Together, these repurposed subunits (with distinct hotspots) form a unique interface that secures the ODC core, priming it for C-terminal tail commitment and underscoring the proteasome’s versatility in substrate recognition.

Another striking discovery is the repurposing of the DUB Rpn11. While canonical substrates engage Rpn11’s Insert-1 β-hairpin to direct chains for deubiquitination and commitment for cleavage, ODC bypasses this site. Instead, its C-terminal tail is first captured by the JAMM motif of Rpn11, which functions as a novel gateway to guide the tail into the AAA+ motor, committing the substrate for degradation. Subsequently, as the tail triggers allosteric activation of proteasome and translocated, it contacts with the H3-helix of Rpn11, which acts as a non-canonical anchor stabilizing the polypeptide (Fig. 2C, E-F, 4B). This dual repurposing of Rpn11, from a DUB to a substrate-tail gate and anchor, highlights the proteasome’s functional adaptability.

### Broader implications for proteostasis and therapeutic development

Our work provides a comprehensive blueprint for direct ubiquitin-independent degradation, expanding our understanding of the proteasome’s functional capabilities. This has profound implications for cellular regulation, particularly in polyamine homeostasis and diseases like cancer, where ODC dysregulation is implicated.^31,32^ Furthermore, by revealing a suite of novel substrate-binding hotspots on the 19S RP, including druggable sites on Rpn10, Rpt5, and Rpn2, our structures provide a roadmap for the design of a new class of therapeutic agents. While current targeted protein degradation (TPD) technologies, such as PROTACs, primarily rely on the ubiquitin system, our findings provide a blueprint for develop ubiquitin-independent degraders that mimic ODC’s engagement strategy. This could revolutionize TPD by providing alternatives for undruggable targets or for therapeutic intervention in cellular contexts where the ubiquitination machinery is compromised.

## Methods

### Express and purification of ODC^ΔC^-Az1^ΔN^ and ODC-Az1

The gene encoding ODC^ΔC^ was constructed to pET28a. Az1^ΔN^ was contructed to pET19b with C-terminal 6×His tag (plasmids were a gift from Nei-li Chan, National Taiwan University, Taipei, China).^35^ Full-length ODC-Az1 expression plasmids were constructed similarly. ODC^ΔC^-Az1^ΔN^ or ODC-Az1 plasmids ware transformed into E. coli BL21 (DE3) cells. Cultures were grown in LB medium with ampicillin and kanamycin to an OD₆₀₀ of 0.6-0.8. Protein expression was induced with 0.5 mM isopropyl β-D-1-thiogalactopyranoside (IPTG) for 18 h at 16°C. Cells were harvested and stored at −80°C.

The purification procedure was identical for both ODC^ΔC^-Az1^ΔN^ and ODC-Az1. The E.coli cell pellet was resuspended in lysis buffer (30 mM HEPES pH 7.5, 5% glycerol, 200 mM NaCl, 5 mM β-mercaptoethanol, 0.5 mM phenylmethanesulfonyl fluoride, and 15 mM imidazole) and disrupted by sonication. The lysate was clarified by centrifugation (18,000 rpm, 1 h, 4°C), the supernatant was applied to a preequilibrated Ni-NTA column (Merck). The column was washed with lysis buffer, and the protein was eluted with elution buffer containing 250 mM imidazole. Eluted fractions were pooled and further purified using a size-exclusion column Superdex 200 10/300 GL (Cytiva) in gel filtration buffer (30 mM HEPES pH 7.5, 250 mM NaCl, 2 mM β-mercaptoethanol, 2 μM pyridoxal 5′-phosphate (PLP)). Peak fractions containing ODC^ΔC^-Az1^ΔN^ or ODC-Az1 complex were further purified using a second size-exclusion column (Hiload 75 16/600, Cytiva). The ODC^ΔC^-Az1^ΔN^ or ODC-Az1 complex was collected and stored in −80℃.

### Purification of WT human 26S proteasome

The Human embryonic kidney (HEK293F) cells were cultured in SMM 293-TII-N medium (SinoBiological) at 37℃ and a 5% CO_2_ atmosphere. Cells from 2 L suspension cultures were harvested by centrifugation, washed with chilled PBS, and stored at −80℃. Cells were homogenized by cryo-milling and dissolved in lysis buffer (50 mM HEPES pH 8.0, 10% glycerol, 10 mM MgCl_2_, 1 mM ATP, 1 mM DTT, 0.1% CA-630). The lysate was clarified by centrifugation (20,000 × g, 60 min), and the supernatant were loaded onto a 50 mL DEAE-Affigel blue column (Bio-Rad) and washed with buffer A (50 mM HEPES pH 8.0, 5% glycerol, 10 mM MgCl_2_, 1 mM ATP, 1 mM DTT). The Proteasome was eluted with a gradient elution from 0%-50% of buffer B (50 mM HEPES pH 8.0, 5% glycerol, 10 mM MgCl_2_, 1 mM ATP, 1 mM DTT with 1 M NaCl). Active fractions, identified by a Suc-LLVY-AMC activity assay, were pooled and loaded onto a Resource Q column (Cytiva). The proteasome was again eluted with a gradient elution from 0%-50% of buffer B, and the fractions were then checked by activity assays conducted either in a 96-well plate or native PAGE. The active fractions eluates were pooled, diluted with Buffer A to reduce the NaCl concentration, and applied to a HiTrap Heparin HP column (Cytiva), from which the proteasome were eluted with a gradient from 20 to 26% gradient of Buffer B. Fractions were further purified by 15%-35% (v/v) glycerol density gradient centrifugation. The purity and integrity of the final proteasome preparation were assessed by native-PAGE and negative-stain EM. Purified proteasomes were concentrated, aliquoted, flash-frozen in liquid nitrogen and stored at −80℃.

### Expression and purification of Rpn11-mutation human 26S proteasome

The gene encoding Rpn11 was amplified by PCR using total cDNAs from HEK293F cell, and constructed into pCDNA3.4 with C-terminal 3 × FLAG tag by ClonExpress II One Step Cloning Kit (Vazyme). Point mutations were introduced using QuikChange site-directed mutagenesis. Plasmids (1 mg) were transfected into 1 L of HEK293F cells (1×10⁶ cells/mL) using polyethyleneimine (PEI). After 48 h, cells were harvested, washed with chilled PBS. Finally, the pellet was stored at −80℃ for further use.

To purify the mutant proteasome, cell pellets were cryo-milled and lysed as described above. The clarified supernatant was incubated with equilibrated Anti-FLAG affinity beads (Smart-Lifesciences, SA042025) for 2 h. Then, the beads were washed with 50 mL of wash buffer (50 mM HEPES pH 8.0, 50 mM NaCl, 10 mM MgCl_2_, 1 mM ATP, 1 mM DTT, 10% glycerol). The complex was eluted with two 15 mL applications of elution buffer containing 0.5 mg/mL 3×FLAG peptide. Subsequently, the eluted samples were further subjected to gel filtration using a Superdex 6 increase 10/300 column (Cytiva) with SEC buffer (50 mM HEPES pH 8.0, 50 mM NaCl, 10 mM MgCl_2_, 1 mM ATP, 1 mM DTT). The purity and integrity of proteasomes were assessed by SDS-PAGE and negative staining, then the final sample were concentrated, aliquoted, flash-frozen in liquid nitrogen and stored at −80℃.

### Native gel-shift and western blotting assay

To detect binding, 26S proteasome was incubated with a 250-fold molar excess of ODC-Az1 in buffer A (25 mM HEPES pH 7.5, 50 mM NaCl, 10% glycerol, 1 mM DTT, 1 mM ATP, 10 mM MgCl₂) for 5 min on ice. Samples were mixed with loading dye and resolved on a 4% native polyacrylamide gel. Proteasome activity was visualized by incubating the gel in buffer containing 100 µM Suc-LLVY-AMC for 10 min at 37℃. For western blotting, the band corresponding to the ODC-Az1-bound proteasome was excised from the native gel. The proteins were analyzed by western blotting and immunoblotted with an anti-ODC/His antibody (Polyclonal).

### Degradation assay of ODC

In vitro degradation assays (40 µL) contained 500 nM purified WT or Rpn11-mutated human 26S proteasome and 500 nM ODC-Az1 in reaction buffer (25 mM HEPES, 50 mM NaCl, 30 mM MgCl₂, 30 mM ATP, 1 mM DTT, 10% glycerol). Reactions were incubated at 37°C for the indicated times and terminated by adding SDS loading buffer. Samples were analyzed by 4-20% SDS-PAGE and western blotting. All assays were performed in triplicate.

### Pull-down assay of Rpn10-ODC

Genes encoding hRpn10 were cloned into pCDNA3.4 vector with a C-terminal HA tag. Genes encoding mNeonGreen2 (mNG2) were cloned into pCDNA3.4 vector with a N-terminal HA tag, and fusion protein was expressed in HEK293F cells. The cells were harvested by centrifugation and stored at −80°C until purification.

The cells were lysed in lysis buffer (50 mM HEPES pH 7.5, 150 mM NaCl, 1 mM DTT, 0.1% CA-630, 10% Glycerol). The lysate was clarified by centrifugation at 20,000 × g for 30 min, and then incubate with Anti-HA Affinity Beads (smart-lifesciences) at 4℃ for 3 h. After washing with wash buffer (50 mM HEPES pH 7.5, 500 mM NaCl, 1 mM DTT, 10% Glycerol), the protein was eluted with elute buffer (50 mM HEPES pH 7.5, 150 mM NaCl, 1 mM DTT, 0.5mg/mL HA peptide, 10% Glycerol). Eluted fraction was further purified by gel-filtration using a Superdex 200 Increase 10/300 GL column (Cytiva). The fractions were assessed by SDS-PAGE, then concentrated, aliquoted, flash frozen in liquid nitrogen and stored at −80℃.

Approximately 200 μg of purified HA-tagged protein (hRpn10-HA or HA-mNG2) was mixed with 600 μg ODC^ΔC^-Az^ΔN^ and then incubated with 20 μL Anti-HA Affinity Beads for 1h at 4℃ with rotation. After incubation, the beads were washed with wash buffer (50 mM HEPES pH7.5, 10 mM MgCl_2_, 150 mM NaCl, 1 mM DTT, 0.1% BSA, 0.02% TritonX-100). Bound proteins were eluted by boiling the beads in 1× SDS loading buffer for western blot analysis.

### Cryo-EM Sample Preparation and Data Collection Sample preparation

For the apo 26S proteasome sample, 2.5 µL of purified proteasome was applied to polylysine-treated, plasma-cleaned grid (R2/1 Cu 200 mesh; Quantifoil). Grids were blotted and plunge-frozen in liquid ethane using a Vitrobot Mark IV (Thermo Fisher Scientific). For the 26S-ODC^ΔC^-Az1^ΔN^ sample, proteasome was incubated with a 50-fold molar excess of ODC^ΔC^-Az1^ΔN^ at 37℃ for 10 min and cross-linked with 0.05% glutaraldehyde for 3-5 min before vitrification. For the 26S-ODC-Az1 sample, proteasome was incubated with a 50-fold molar excess of ODC-Az1 at 10℃ for 2-6 min before vitrification. Grids were stored in liquid nitrogen.

### Data collection

Cryo-EM movies were collected on a Titan Krios electron microscope (Thermo Fisher Scientific) equipped with a Cs corrector, operating at 300 kV. Data were collected automatically using EPU v2.11 with a defocus range of −0.8 to −2.5 µm. Movies were recorded on a K3 Summit direct electron detector (Gatan) with a total dose of 50 e⁻/Å². For the 26S-ODC^ΔC^-Az1^ΔN^ sample, movies were recorded in counting mode (1.1 Å/pixel). For the 26S alone and 26S-ODC-Az1 samples, movies were recorded in super-resolution mode and binned by 2 (final pixel sizes of 1.0842 Å and 1.1 Å, respectively).

### Cryo-EM data processing

Cryo-EM single-particle analysis was performed using Relion3.1^68,69^ and cryoSPARC4.6.^70^ Movies were motion-corrected using MotionCorr2^71^, and CTF parameters were determined using CTFFIND4.^72^

For the free 26S proteasome dataset, automated particle picking using relion from 4,380 micrographs followed by reference-free 2D classification yielded 741,026 particles. After two rounds of 3D classification (during which the opposing 19S cap was extracted and combined), CTF refinement and Bayesian polishing in RELION, the particles were transferred to cryoSPARC for further 3D classification. The final set of 110,674 particles was refined to a consensus map at 3.4 Å resolution. Subsequent local refinements of the 19S and 20S regions yielded maps at 3.5 Å and 3.2 Å resolution, respectively. A similar procedure was used to process the other datasets.

For the 26S-ODCΔC-Az1ΔN dataset, from approximately 30,000 micrographs, after pre-processing, 1,435,434 particles remained. Subsequent 3D classification generated ten distinct classes, of which the fifth and tenth exhibiting additional density were selected for further processing. The tenth class underwent CTF refinement, Bayesian polishing, and another round of 3D classification, yielding a subset of 73,575 particles. Homogeneous refinement of this subset produced a consensus map at 3.7 Å resolution, designated as the R3 state. Focused local refinements of the base, lid, and 20S regions achieve final resolutions of 4.1 Å, 4.5 Å, and 3.5 Å, respectively. Similarly, processing of the fifth class yielded two final subsets corresponding to the R1 state (13,668 particles) and the R2 state (12,670 particles) states, which were refined to overall resolutions of 4.4 Å and 4.5 Å, respectively.

For the 26S-ODC-Az1 dataset, pre-processing of 27,500 micrographs yielded 4,363,576 particles. Subsequent 3D classification identified two major populations: a resting (1,076,303) and an activated (1,006,343) states. The resting-state particles were further processed using 3D variability analysis (3DVA) and multiple rounds of focused 3D classification on the ODC-Az1 and Rpn2-Rpn10-Rpt4/5 coiled-coil regions. This yielded three final subsets corresponding to the B1 (43,171 particles), B2 (30,180 particles), and B3 (11,467 particles) states, which were refined to resolutions of 3.4 Å, 3.6 Å, and 3.9 Å, respectively. Subsequent local refinements were performed on the 19S and 20S regions to improve the resolvability especially for the 19S RP, then the two portions were merged to generate final composite maps. For the activated-state population, a similar processing strategy, which included focused 3D classification centered on the ODC-Az1 region, yielded five final subsets corresponding to the A1 (27,448 particles), A2 (57,780 particles), A3 (56,498 particles), A4 (61,424 particles), and A5 (68,388 particles) states, refined to resolutions ranging from 3.4 Å to 4.4 Å.

The overall resolutions for all of the cryo-EM maps were determined based on the gold-standard criterion using a Fourier shell correlation (FSC) of 0.143. All maps were sharpened using DeepEMhancer 0.14^73^, and composite maps were generated using the “vop maximum” command in ChimeraX 1.8.^74–76^

### Atomic model building and validation

For the resting state, initial models were generated based on existing structures of the resting-state human 26S proteasome (5VSF)^49^ and ODC^ΔC^-Az1^ΔN^ crystal structure (4ZGY)^35^, along with AlphaFold3^77^ predicted models for SEM1. We first rigid-body fitted the models into the corresponding cryo-EM maps using Chimera.^78^ The models then underwent flexible fitting against the corresponding map using Rosetta^79^ real-space refinement package, followed by several rounds of refinement using the Real_space_refine module in PHENIX^80^ to optimize the models. A similar procedure was followed for the activated state, using initial models based on the activated stated human 26S proteasome structure (PDB: 6MSE,6MSK).^18^

UCSF Chimera^78^ and ChimeraX^74–76^ were applied for figure generation, distance/rotation measurements, and surface potential analysis.

### BS3 Mediated Crosslinking Mass Spectrometry Analysis

The proteins were precipitated and digested for 16 h at 37℃ by trypsin at an enzyme-to-substrate ratio of 1:50 (w/w). The tryptic digested peptides were desalted and loaded on an in-house packed capillary reverse-phase C18 column (30cm length, 75 µM ID x 360 µM OD, ReproSil-Pur C18-AQ 1.9 uM resin, 120 Å pore diameter, Dr. Maisch GmbH) connected to an Easy LC 1200 system. The analytical column temperature was set at 55 ℃ during the experiments. The mobile phase and elution gradient used for peptide separation were as follows: 0.1% formic acid in water as buffer A and 0.1% formic acid in 80% acetonitrile as buffer B, 0-1 min, 5%-6% B; 1-96 min, 6-36% B; 96-107 min, 36%-60% B, 107-108 min, 60%-100% B, 108-120 min, 100% B. The flow rate was set as 300 nL/min. Data-dependent MS/MS analysis was performed with a Q Exactive Orbitrap mass spectrometer (Thermo Scientific). Peptides eluted from the LC column were directly electrosprayed into the mass spectrometer with the application of a distal 2.5-kV spray voltage. A cycle of one full-scan MS spectrum (m/z 300-1800) was acquired followed by top 20 MS/MS events, sequentially generated on the first to the twentith most intense ions selected from the full MS spectrum at a 28% normalized collision energy. Full scan resolution was set to 70,000 with automated gain control (AGC) target of 3e6. MS/MS scan resolution was set to 17,500 with isolation window of 1.8 m/z and AGC target of 1e5. The number of microscans was one for both MS and MS/MS scans and the maximum ion injection time was 50 and 100 ms, respectively. The dynamic exclusion settings used were as follows: charge exclusion, 1 and >8; exclude isotopes, on. MS scan functions and LC solvent gradients were controlled by the Xcalibur data system (Thermo Scientific).

Cross-linked peptides were identified using pLink2 software (version 2.3.9, pFind Team, Beijing, China) as described previously.^81^ Search parameters in pLink were: enzyme: trypsin; missed cleavages: 3; precursor and fragment tolerance: 20 ppm. Oxidation of methionine was set as variable modification. The results were filtered by applying a 5% FDR cutoff at the spectral level.

The proteasome-ODC complex was cross-linked with BS3, then precipitated and digested with trypsin. Tryptic peptides were analyzed by LC-MS/MS on a Q Exactive Orbitrap mass spectrometer (Thermo Scientific). Cross-linked peptides were identified using pLink2 software with a 5% false discovery rate cutoff.

## Reporting summary

### Data availability

The cryo-EM maps determined for the 26S-ODC-Az1 have been deposited at the Electron Microscopy Data Bank with the accession codes ???, ???, ???, ???, ???, ???, ???, ???, ???, ???, ???, ???, and the associated atomic models have been deposited in the PDB with the accession codes ???, ???, ???, ???, ???, ???, ???, ???, ???, ???, ???, ???. For gel source images, see Supplementary.

## Acknowledgements

We thank Nei-Li Chan for the plasmids expressing ODC^ΔC^ and Az1^ΔN^. We thank the staff at the Large-scale Protein Preparation System, Electron Microscopy System, Database and Computation System at the National Facility for Protein Science in Shanghai (NFPS), for instrument support and technical assistance. This work was supported by grants from the Strategic Priority Research Program of CAS (XDB37040103, XDB0570303), the NSFC (32130056, 31872714 to Y.C.), the National Key R&D Program of China (2024YFA1803102), Shanghai Pilot Program for Basic Research from CAS (ICYJ-SHFY-2022-008 to Y.C.).

## Author contributions

K.C. and Y.C. conceived and designed the experiments. K.C., Y.W., X.Y., Xuyang Yuan, and Z.D. purified proteins. K.C., Y.W., X.Y., Xuyang Yuan, and C.X. performed functional analysis. K.C. and Y.Y. performed MS analysis. K.C. and C.D. performed data collection. K.C. reconstructed the structures. K.C. and Y.C. analyzed the structures. K.C., Y.W., Y.Y. and Y.C. wrote the manuscript.

## Supplementary information

**Fig. S1.**
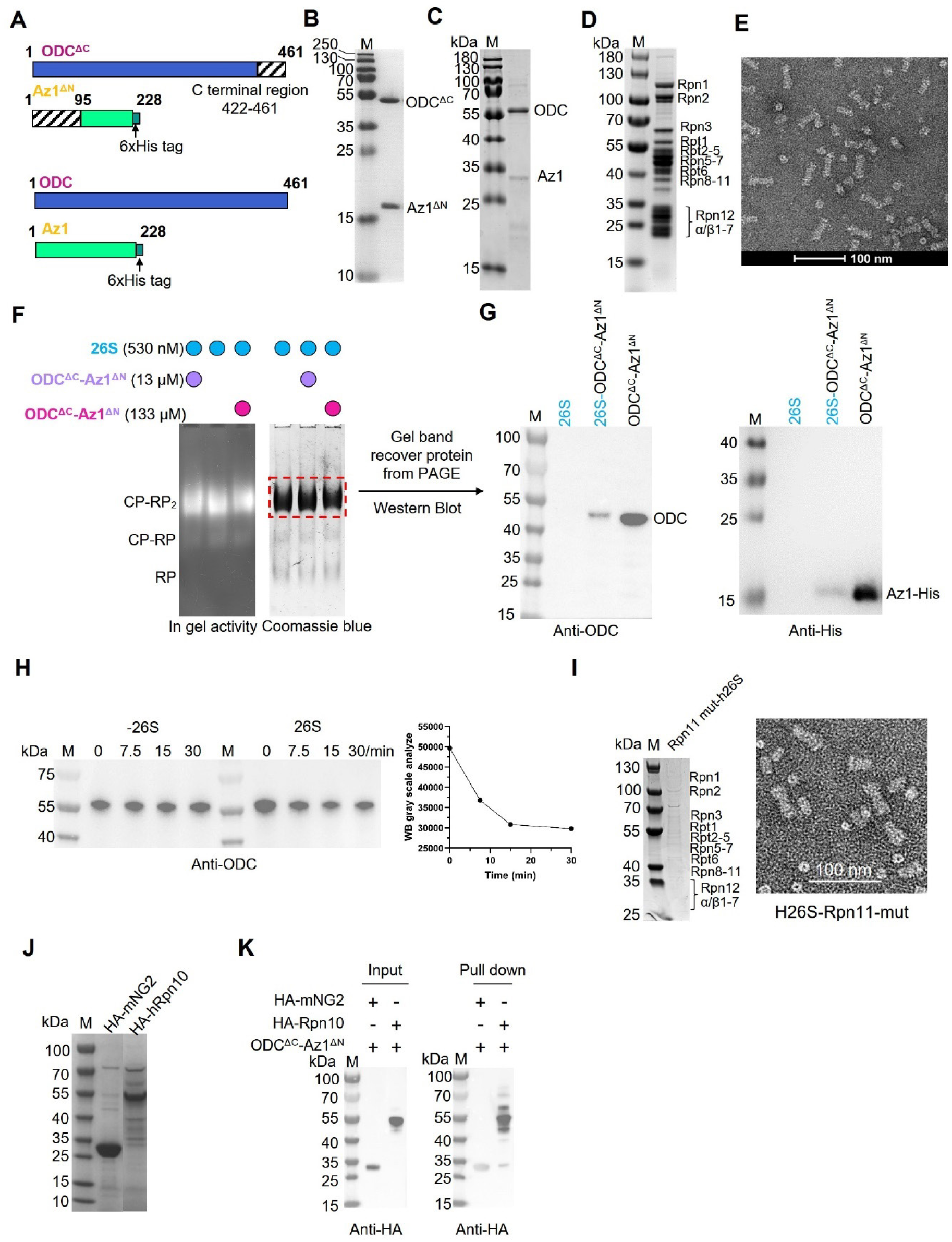
Biochemical characterization of ODC-Az1 complexes and their interaction with the human 26S proteasome. (A) Schematic diagrams of ODC^ΔC^-Az1^ΔN^ (up) and full-length ODC-Az1 (down). (B, C) SDS-PAGE analysis showing the purified ODC^ΔC^-Az1^ΔN^ (B) and ODC-Az1 (C) complexes. (D, E) SDS-PAGE analysis of the purified human 26S proteasome (D), and representative negative staining EM (NS-EM) of the proteasome (E). (F) Native gel shift assay demonstrating the association between the 26S proteasome and ODC^ΔC^-Az1^ΔN^. The complex is visualized by an in-gel activity assay (left) and Coomassie brilliant blue staining (right). (G) Western blot analysis of the shifted band from the native gel, confirming the presence of ODC and Az1. (H) *In vitro* degradation assay showing that purified proteasomes degrade full-length ODC-Az1, monitored by western blot. (I) SDS-PAGE analysis of the purified Rpn11-mutant 26S proteasome. The sample was also checked by NS-EM (right). (J) SDS-PAGE analysis of purified HA-tagged hRpn10 and mNG2 used in pull-down assays. (K) Pull-down assay showing that HA-Rpn10, but not the HA-mNG2 negative control, interacts with ODC^ΔC^-Az1^ΔN^.

**Fig. S2.**
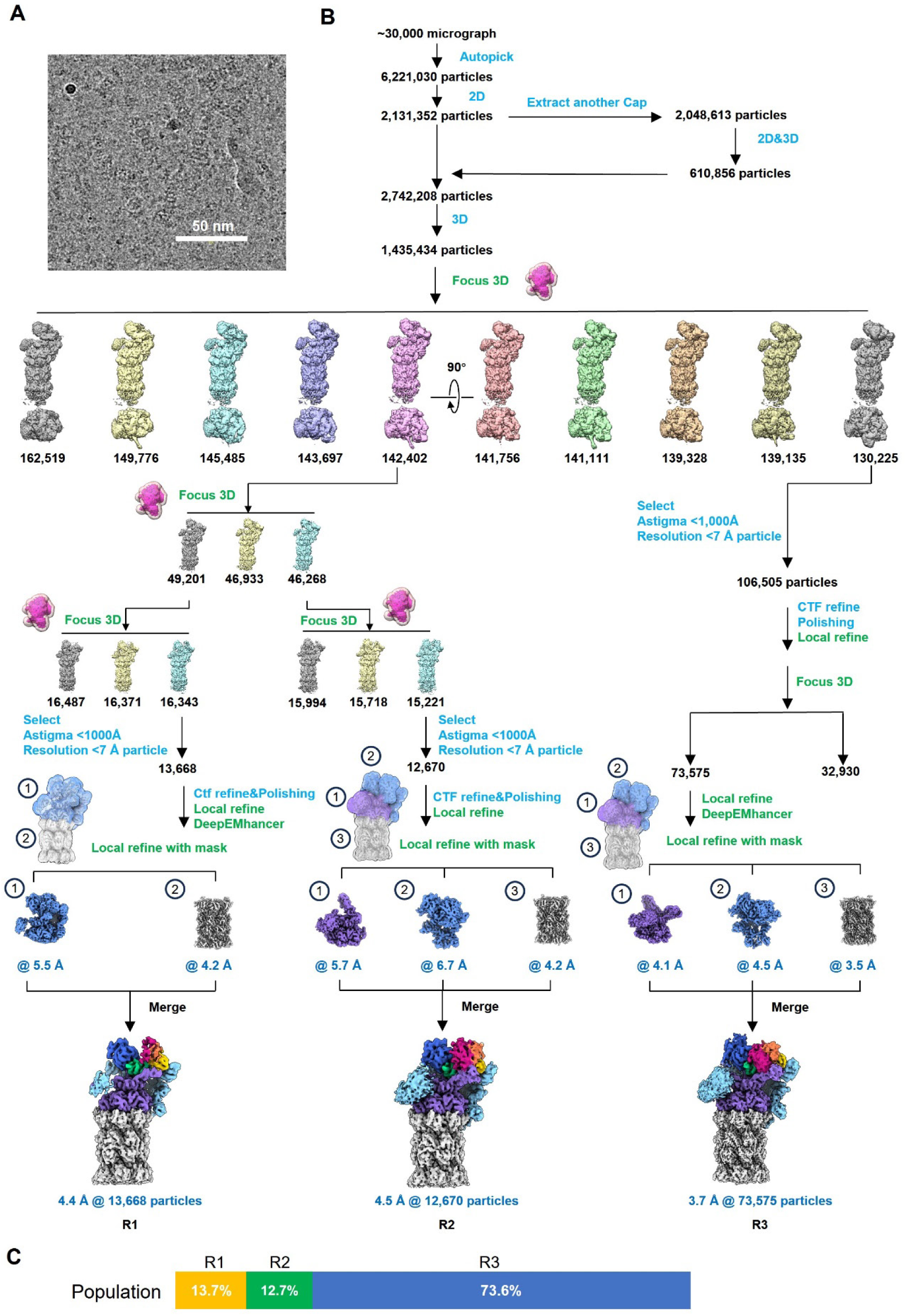
Cryo-EM data processing procedure for the 26S-ODC^ΔC^-Az1^ΔN^ complex. (A) A representative cryo-EM micrograph. (B) Data processing workflow. (C) Particle populations among the R1, R2 and R3 states.

**Fig. S3.**
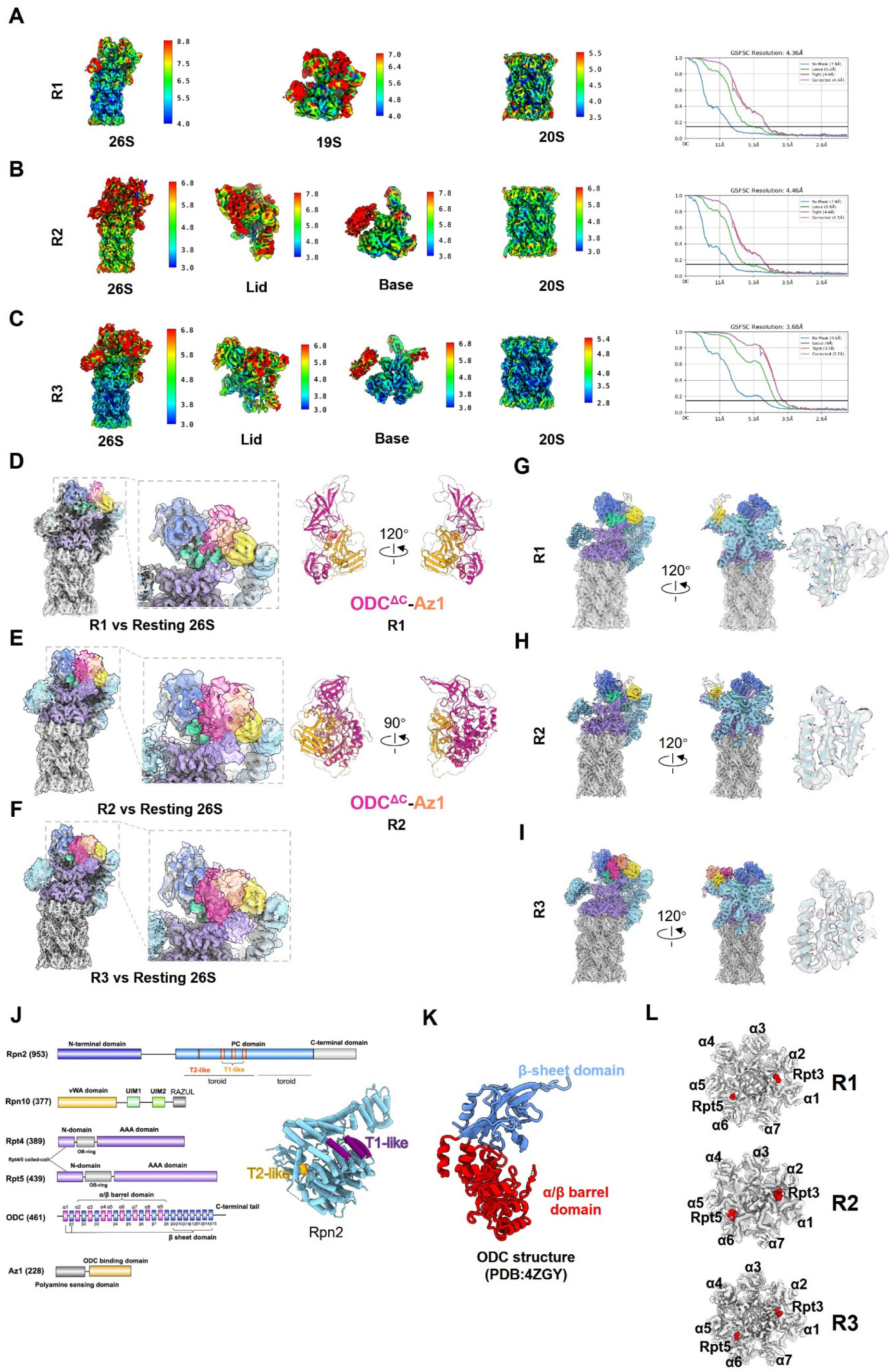
Cryo-EM map and model quality for the 26S-ODC^ΔC^-Az1^ΔN^ complex. (A-C) Local resolution evaluation and Fourier shell correlation (FSC) curves for the R1 (A), R2 (B), and R3 (C) states, including focused refined portions of the map. (D-F) Comparison of the R1 (D), R2 (E), and R3 (F) maps with the apo 26S proteasome map, showing the location of the extra density for ODC^ΔC^-Az1^ΔN^. Also showed is the fit of the ODC^ΔC^-Az1^ΔN^ crystal structure (PDB:4ZGY) into the extra density of the R1 (D) and R2 (E) states (transparent surface). (G-I) Model-map fitting for R1 (G), R2 (H), and R3 state (I), and a representative high-resolution structure feature for each map. (J) Domain diagram of ODC, Az1, and key contacting 19S subunits. The T1-/T2-like sites of Rpn2 PC domain are also illustrated in the model. (K) Domain organization of ODC. (L) Top view of the 20S CP, showing the closed-gate status in the resting R1, R2 and R3 states.

**Fig. S4.**
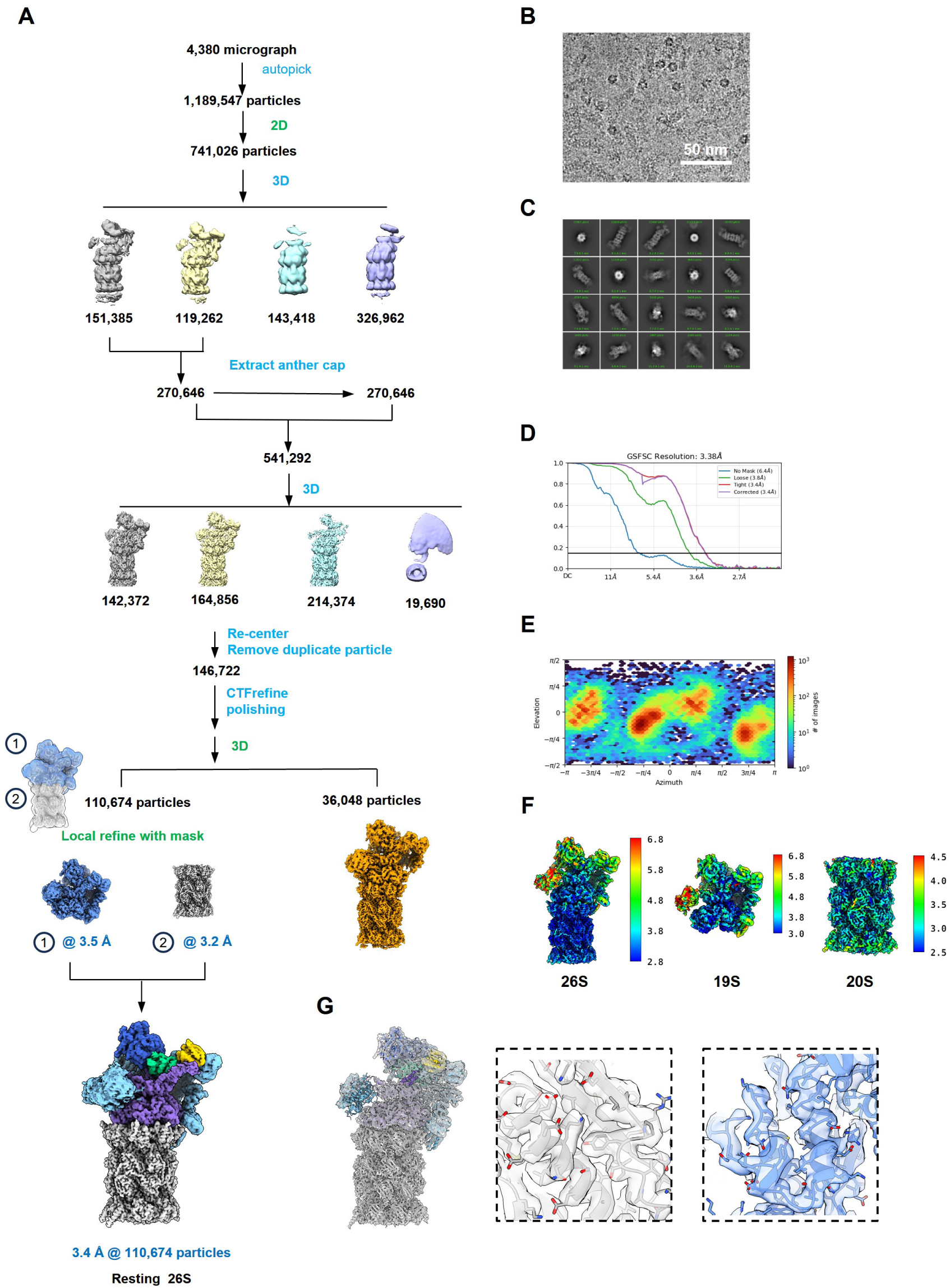
Cryo-EM analysis of the free 26S proteasome. (A) Data processing workflow for the human 26S proteasome alone. (B) A representative micrograph. (C) Reference-free 2D class averages. (D) Gold-standard FSC curves for the consensus and focuse-refined maps. (E) Angular distribution of particles in the final reconstruction. (F) Local resolution evaluation of maps. (G) Model-map fitting for 26S proteasome.

**Fig. S5.**
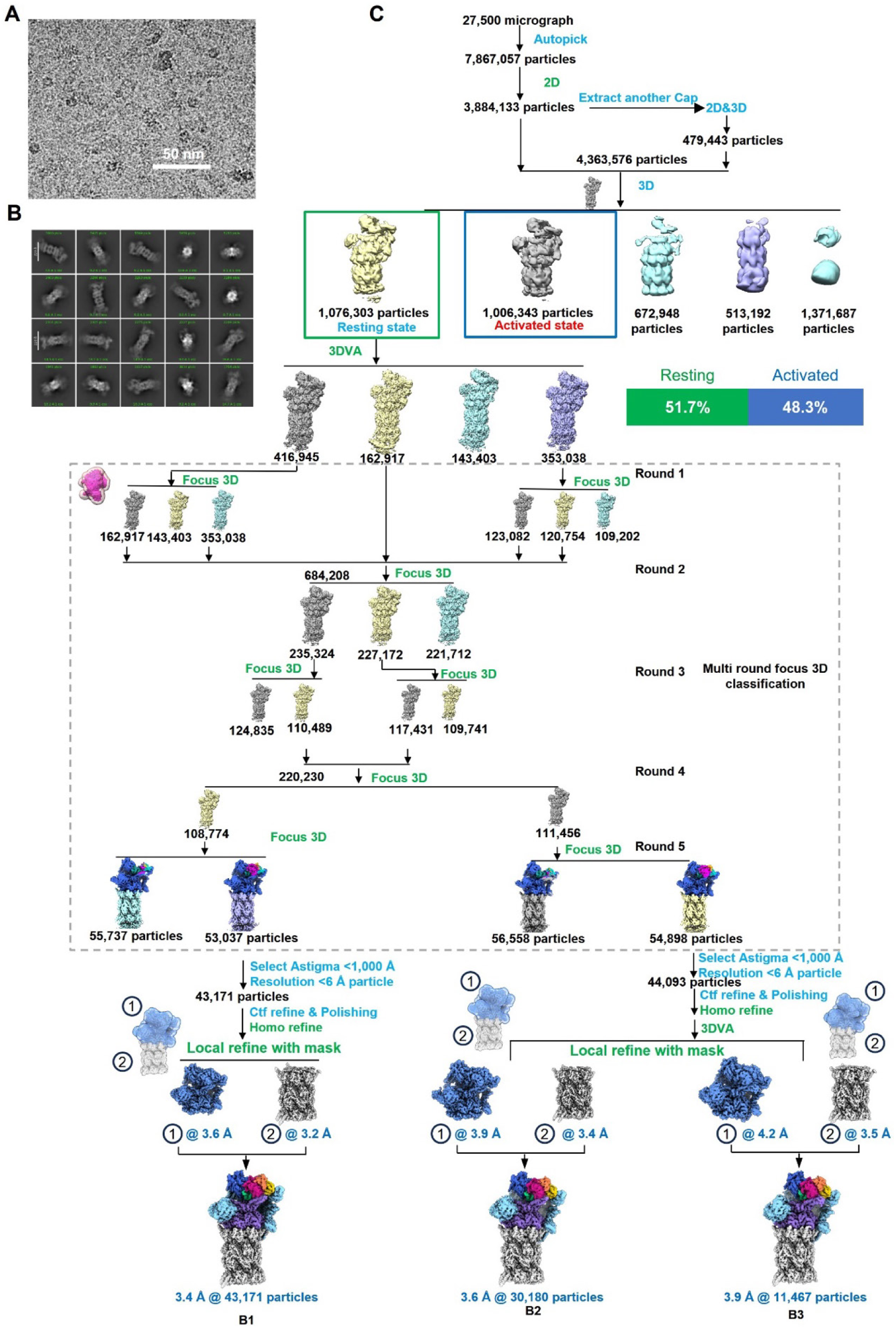
Cryo-EM data processing of the 26S-ODC-Az1 complex, especially for the resting state. (A) A representative micrograph. (B) Reference-free 2D class averages. (C) Overall data processing workflow, showing the initial classification into resting and activated populations.

**Fig. S6.**
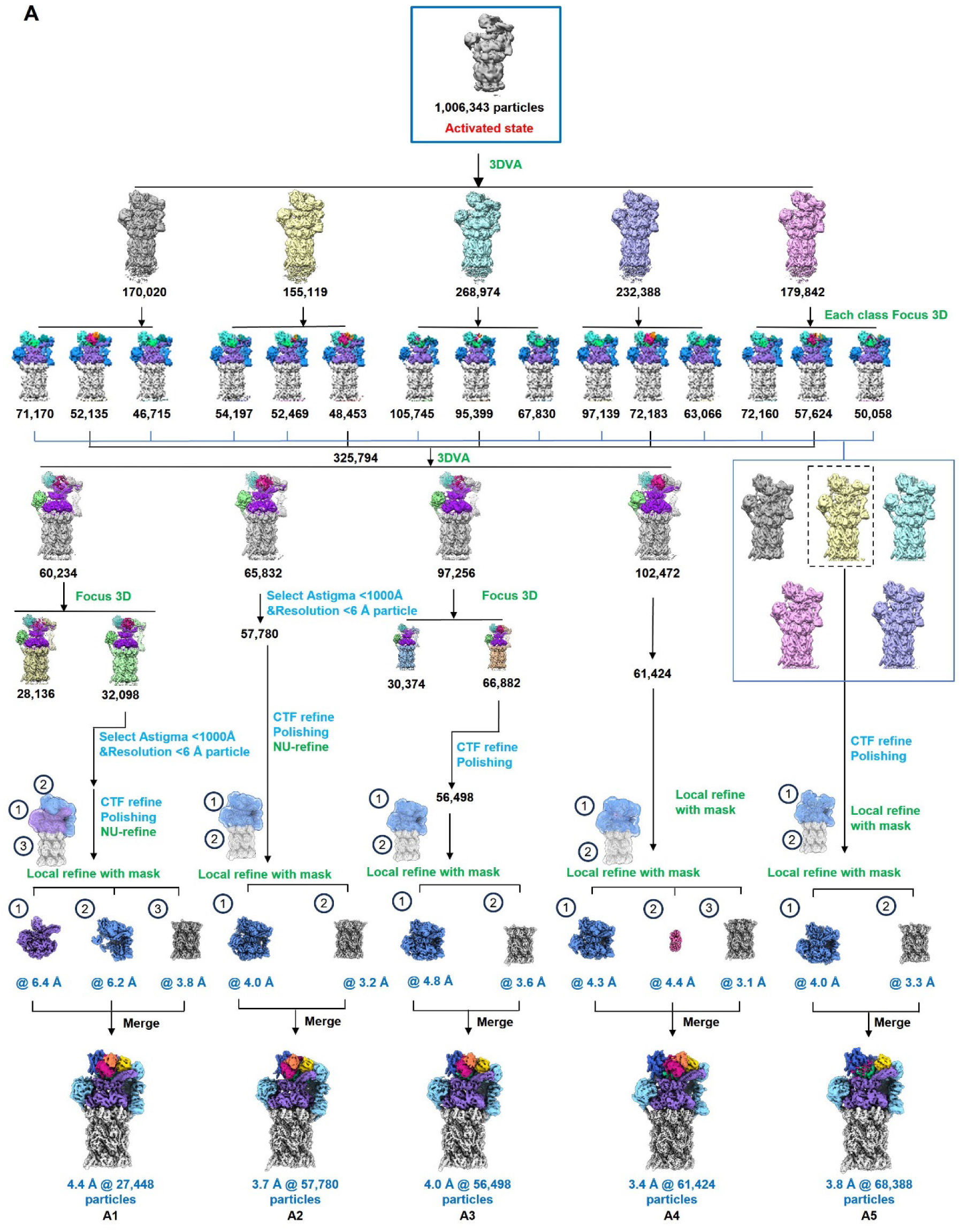
Cryo-EM data processing of the 26S-ODC-Az1 complex for the activated state population.

**Fig. S7.**
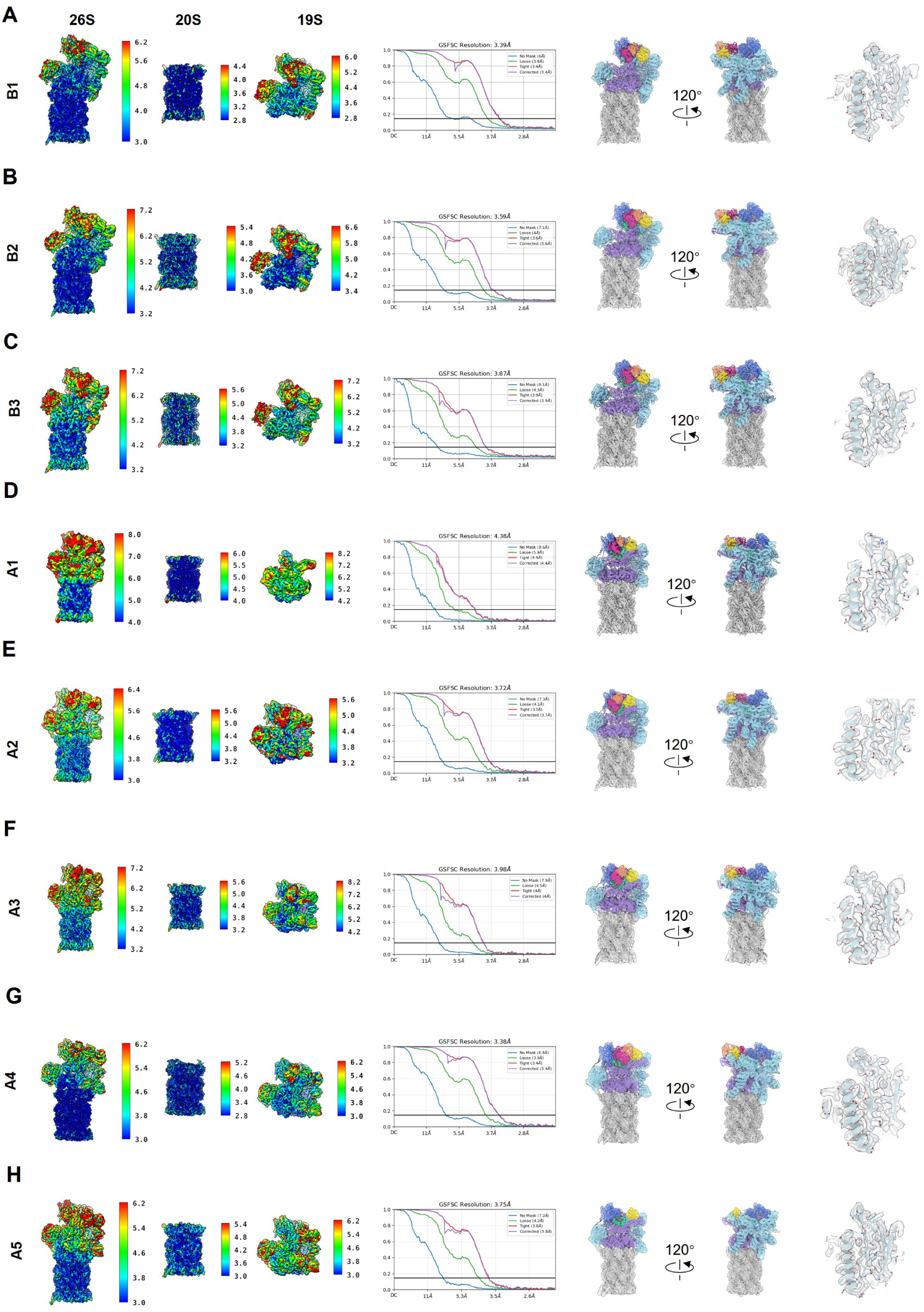
Cryo-EM map and model quality evaluations for the 26S-ODC-Az1 complex. (A-H) For each of the B1-B3 and A1-A5 states, panels show local resolution maps, gold-standard FSC curves, map-model fitting, and high-reslution structure features for the final reconstructions: B1 (A), B2 (B), B3 (C), A1 (D), A2 (E), A3 (F), A4 (G), and A5 (H).

**Fig. S8.**
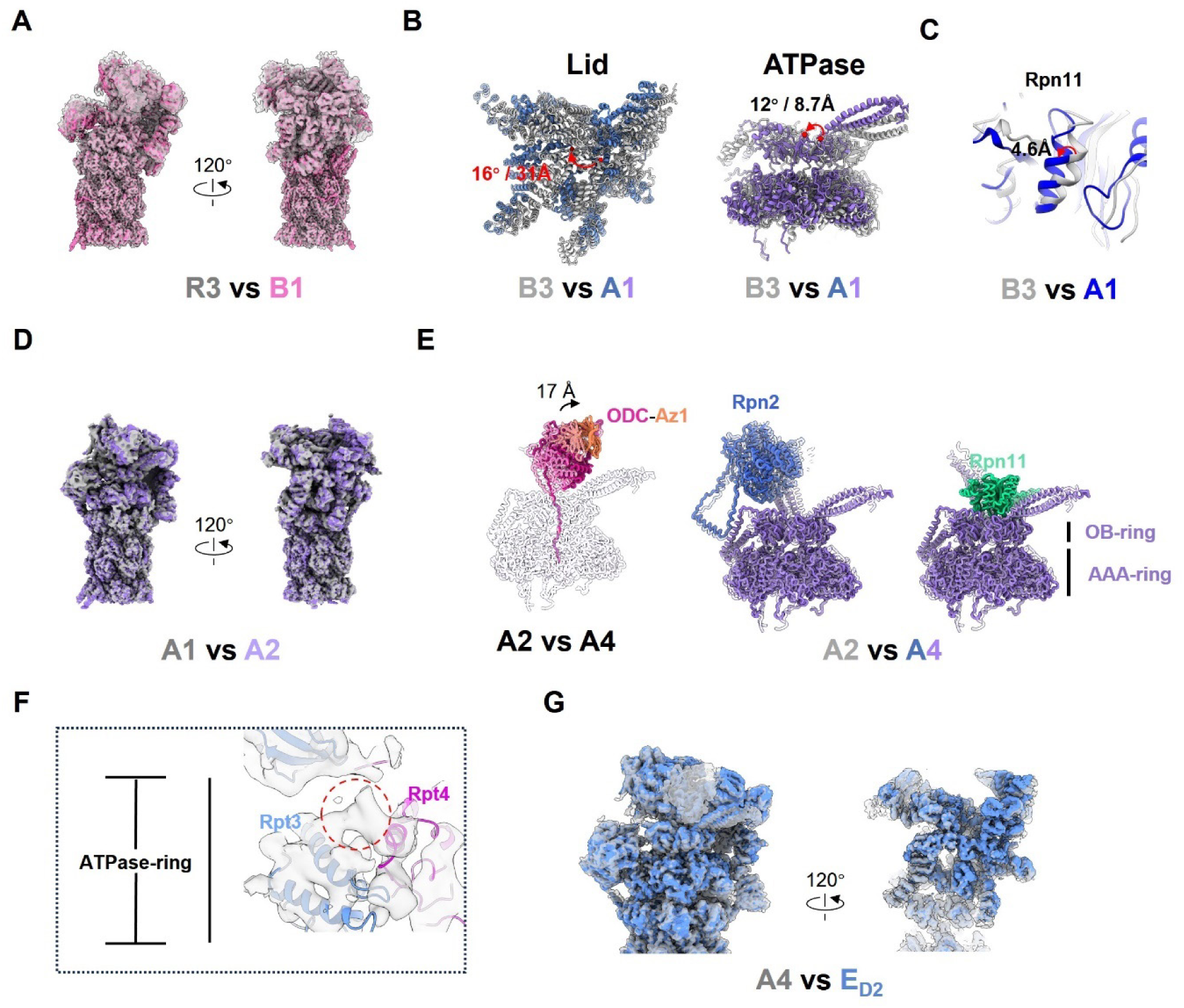
Conformational changes of the 26S proteasome induced by ODC binding and C-terminal tail induced proteasome activation. (A) Superposition of the R3 (grey) and B1 (light pink) cryo-EM maps. (B) Structural comparison of the B3 (grey) and A1 (blue for lid; purple for ATPase ring) states. (C) Conformational variation in Rpn11 subunit between B3 (gray) and A1 (blue) state, showing a 4.6 Å shift of the H3 helix of Rpn11. (D) Superposition of the A1 (gray) and A2 (purple) maps. (E) Structural comparisons between between A2 (transparent) and A4 (color) states, illustrating the movement of ODC-Az1 and the key subunits. Conformational variation between B3(gray) and A1(blue) of selected subunits (shift distance labeled). (F) The extra density (grey) corresponding to the ODC C-terminal tail making initial contact with the Rpt3/4 pore-1 loops in the A1 state. Rpt3 model was color in cornflower blue, Rpt4 model in Magenta. (G) Comparison of cryo-EM maps A4 with a canonical ubiquitin-substrate-engaged E_D2_ (EMD: 9222) state.

**Fig. S9.**
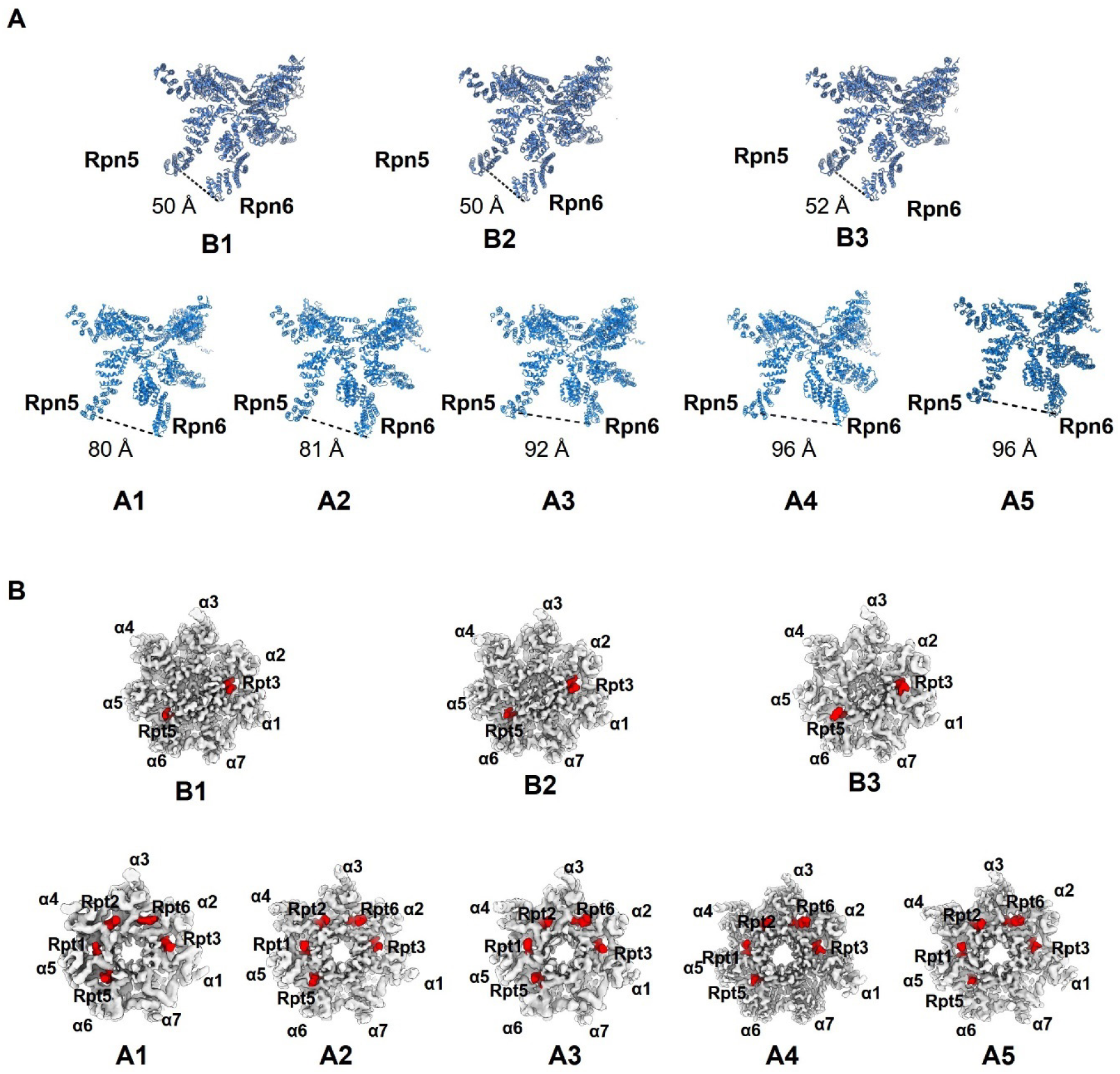
Indications of proteasome activation of the 26S-ODC-Az1 complex. (A) Measurement of the distance between Rpn5 and Rpn6 in the resting B1-B3 states (up) and activated A1-A5 states (down), showing a progressive increase upon activation. (B) Top views of the 20S CP gate, showing it is closed in all B1-B3 states (up) and open in all A1-A5 states (down).

**Fig. S10.**
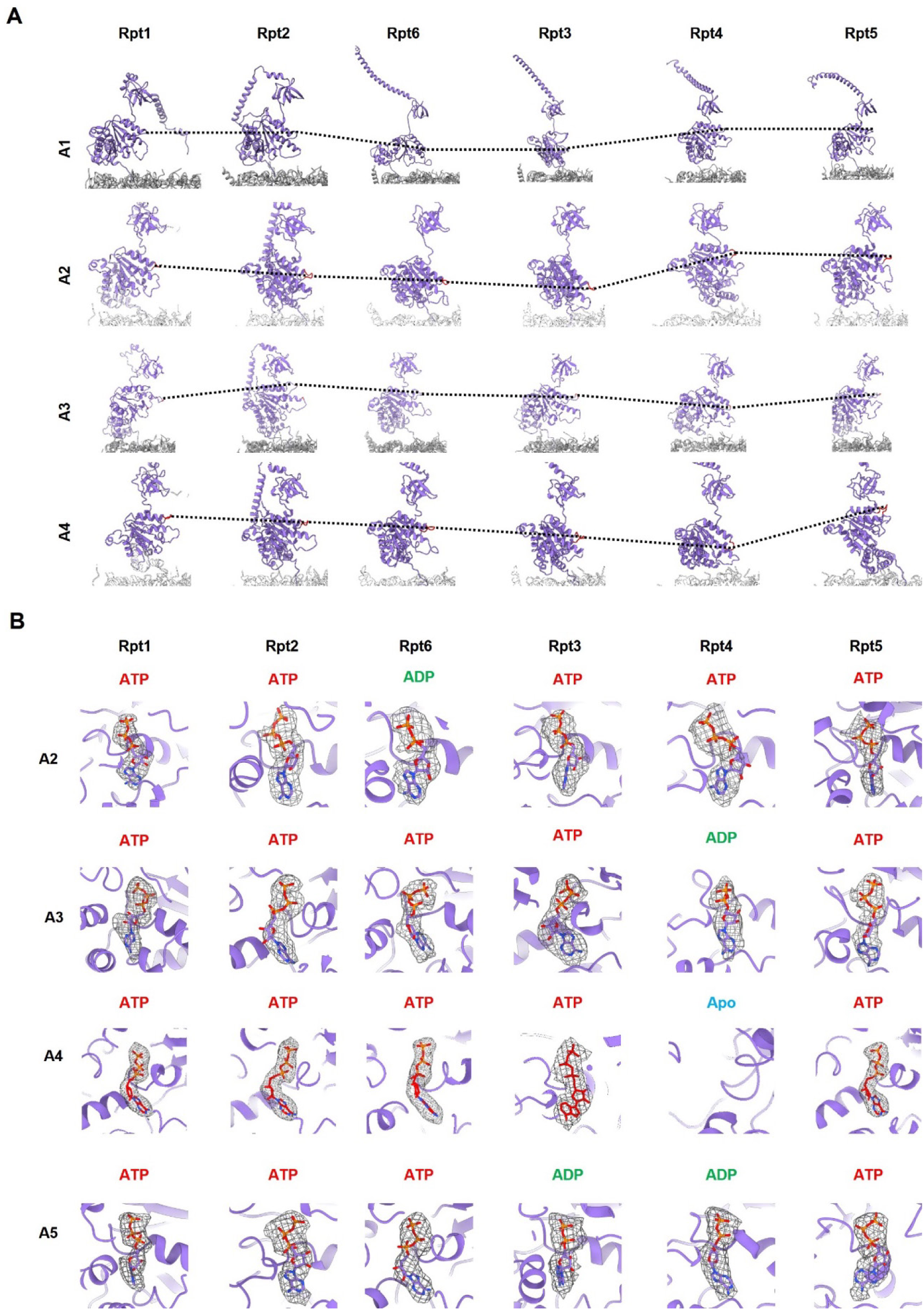
ATPase ring configuration during translocation. (A) Configuration of the Rpt pore-1 loops in the ATPase ring in A1-A4 states. (B) Nucleotide status in the Rpt subunits across the better resolved acrivated A2-A5 states.

**Table S1.**
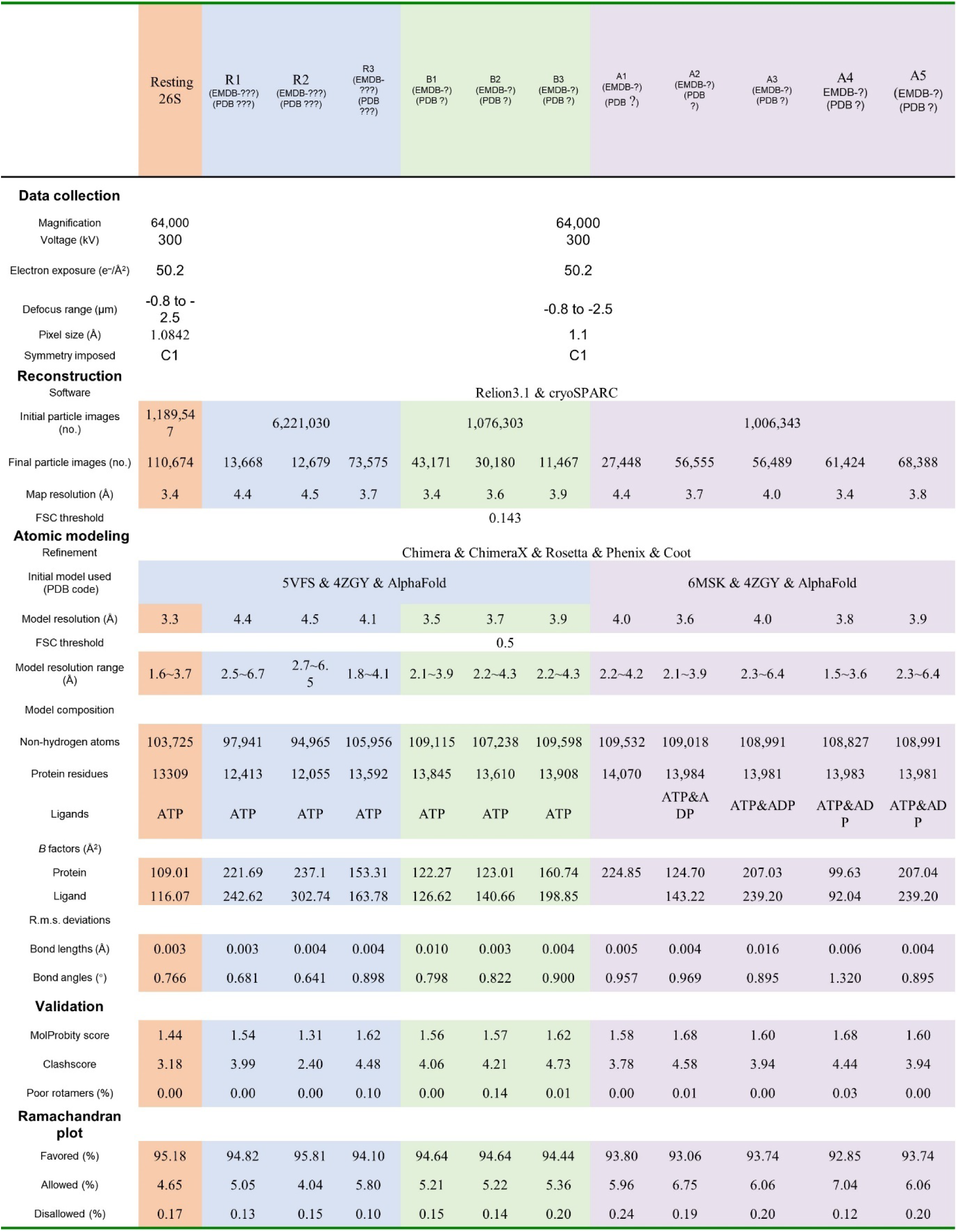
Cryo-EM data collection and refinement statistics.

**Table S2.** IP-ID MS of 26S-ODC^ΔC^-Az1^ΔN^ in native gel.

| Protein | Coverage [%] | # PSMs | # Unique Peptides |
| --- | --- | --- | --- |
| Rpn3 | 72 | 165 | 45 |
| Rpt1 | 76 | 162 | 39 |
| Rpn7 | 75 | 143 | 40 |
| Rpn6 | 68 | 156 | 34 |
| Rpn9 | 74 | 102 | 29 |
| Rpt4 | 70 | 136 | 25 |
| Rpn11 | 87 | 100 | 18 |
| α6 | 79 | 111 | 25 |
| Rpn8 | 62 | 71 | 16 |
| β6 | 75 | 66 | 18 |
| α4 | 64 | 67 | 18 |
| α2 | 57 | 57 | 12 |
| α5 | 63 | 60 | 15 |
| β5 | 65 | 61 | 17 |
| α1 | 71 | 78 | 18 |
| β2 | 71 | 44 | 14 |
| β3 | 51 | 58 | 15 |
| α3 | 77 | 52 | 15 |
| UCHL5 | 78 | 56 | 21 |
| β4 | 90 | 63 | 19 |
| Rpn12 | 52 | 54 | 17 |
| β7 | 48 | 49 | 9 |
| Rpn10 | 44 | 45 | 15 |
| ODC | 33 | 44 | 18 |
| α7 | 41 | 51 | 13 |
| β1 | 69 | 27 | 10 |
| Az1 | 43 | 20 | 8 |

**Table S3.**
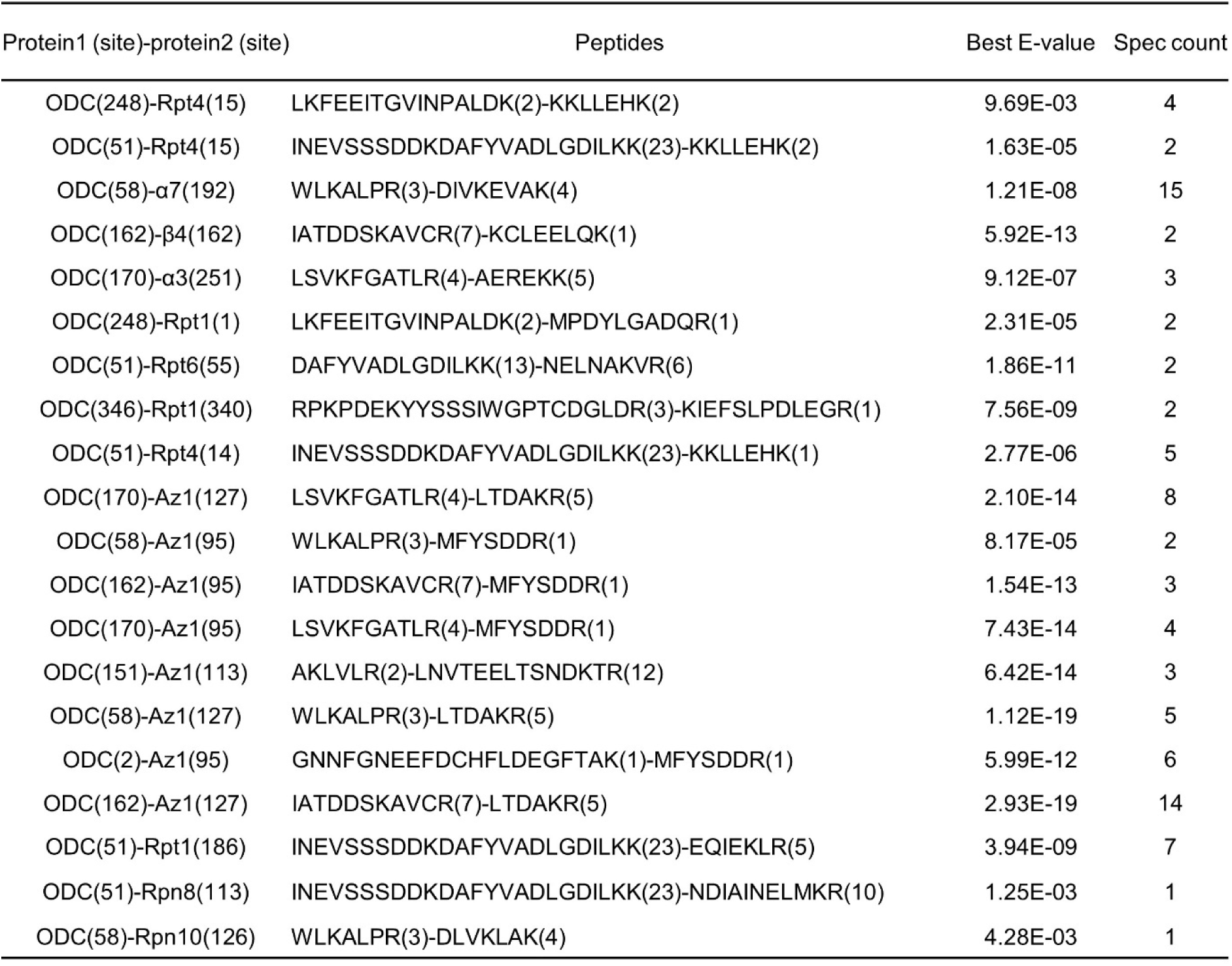
XL-MS of 26S-ODC^ΔC^-Az1^ΔN^.

| Protein1 (site)-protein2 (site) | Peptides | Best E-value | Spec count |
| --- | --- | --- | --- |
| ODC(248)-Rpt4(15) | LKFEEITGVINPALDK(2)-KKLLEHK(2) | 9.69E-03 | 4 |
| ODC(51)-Rpt4(15) | INEVSSSDDKDAFYVADLGDILKK(23)-KKLLEHK(2) | 1.63E-05 | 2 |
| ODC(58)-α7(192) | WLKALPR(3)-DIVKEVAK(4) | 1.21E-08 | 15 |
| ODC(162)-β4(162) | IATDDSKAVCR(7)-KCLEELQK(1) | 5.92E-13 | 2 |
| ODC(170)-α3(251) | LSVKFGATLR(4)-AEREKK(5) | 9.12E-07 | 3 |
| ODC(248)-Rpt1(1) | LKFEEITGVINPALDK(2)-MPDYLGADQR(1) | 2.31E-05 | 2 |
| ODC(51)-Rpt6(55) | DAFYVADLGDILKK(13)-NELNAKVR(6) | 1.86E-11 | 2 |
| ODC(346)-Rpt1(340) | RPKPDEKYYSSSIWGPTCDGLDR(3)-KIEFSLPDLEGR(1) | 7.56E-09 | 2 |
| ODC(51)-Rpt4(14) | INEVSSSDDKDAFYVADLGDILKK(23)-KKLLEHK(1) | 2.77E-06 | 5 |
| ODC(170)-Az1(127) | LSVKFGATLR(4)-LTDAKR(5) | 2.10E-14 | 8 |
| ODC(58)-Az1(95) | WLKALPR(3)-MFYSDDR(1) | 8.17E-05 | 2 |
| ODC(162)-Az1(95) | IATDDSKAVCR(7)-MFYSDDR(1) | 1.54E-13 | 3 |
| ODC(170)-Az1(95) | LSVKFGATLR(4)-MFYSDDR(1) | 7.43E-14 | 4 |
| ODC(151)-Az1(113) | AKLVLR(2)-LNVTEELTSNDKTR(12) | 6.42E-14 | 3 |
| ODC(58)-Az1(127) | WLKALPR(3)-LTDAKR(5) | 1.12E-19 | 5 |
| ODC(2)-Az1(95) | GNNFGNEEFDCHFLDEGFTAK(1)-MFYSDDR(1) | 5.99E-12 | 6 |
| ODC(162)-Az1(127) | IATDDSKAVCR(7)-LTDAKR(5) | 2.93E-19 | 14 |
| ODC(51)-Rpt1(186) | INEVSSSDDKDAFYVADLGDILKK(23)-EQIEKLR(5) | 3.94E-09 | 7 |
| ODC(51)-Rpn8(113) | INEVSSSDDKDAFYVADLGDILKK(23)-NDIAINELMKR(10) | 1.25E-03 | 1 |
| ODC(58)-Rpn10(126) | WLKALPR(3)-DLVKLAK(4) | 4.28E-03 | 1 |

**Table S4.**
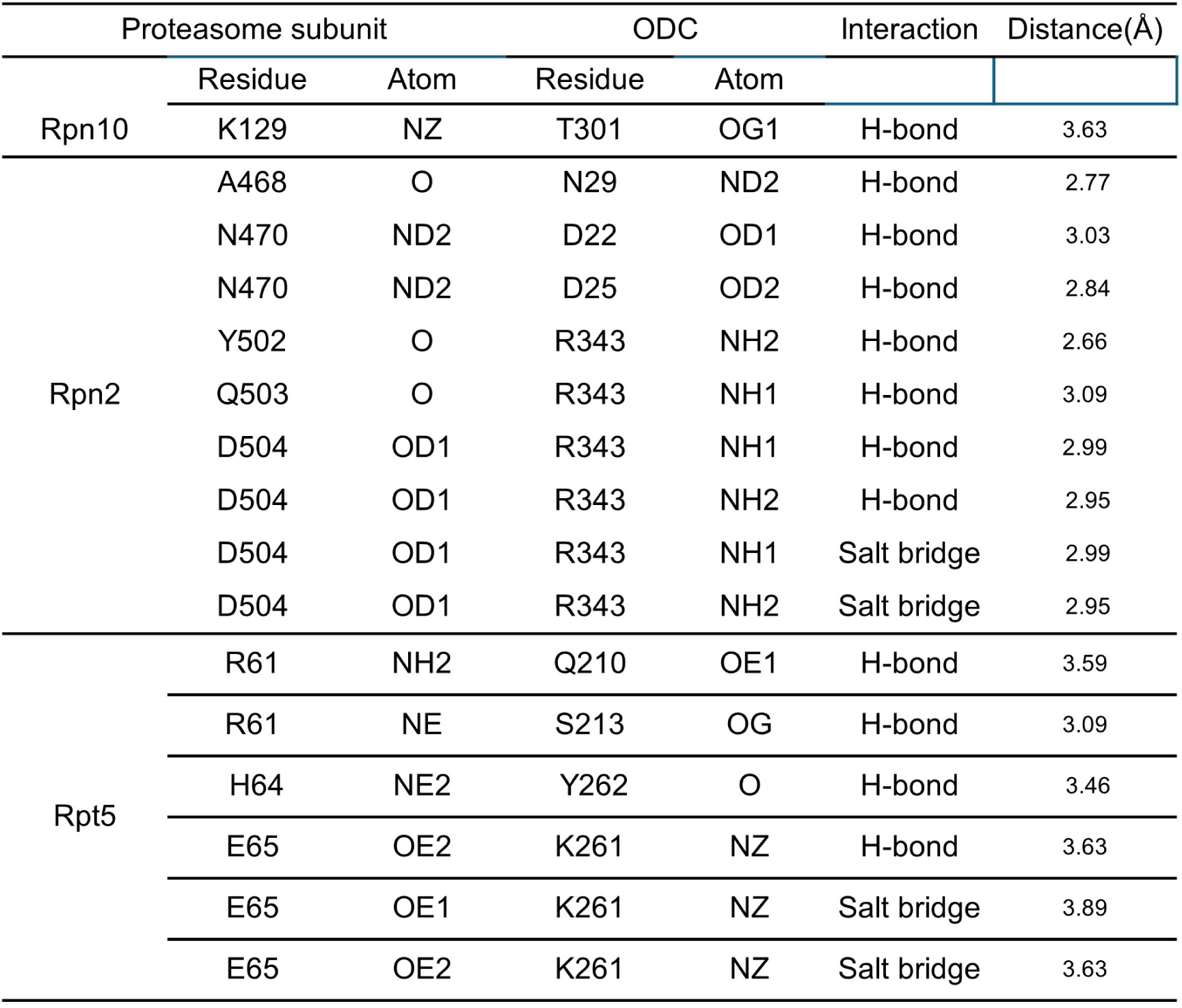
Interaction network between 26S proteasome and ODC^ΔC^ analyzed by PISA.

**Table S5.** Interaction network between 26S proteasome and ODC in B1 state analyzed by PISA.

|  | Proteasome subunit |  | ODC |  | Interaction | Distance(Å) |
| --- | --- | --- | --- | --- | --- | --- |
|  | Residue | Atom | Residue | Atom |  |  |
| Rpn10 | K106 | NZ | E368 | OE2 | H-bond | 2.82 |
|  | K106 | NZ | E368 | OE1 | H-bond | 3.05 |
|  | K129 | NZ | S303 | OG | H-bond | 3.11 |
|  | R130 | NH1 | D305 | OD2 | H-bond | 2.77 |
|  | R130 | NH1 | D305 | OD1 | H-bond | 3.11 |
|  | K133 | NZ | D304 | OD2 | H-bond | 2.40 |
|  | R42 | O | R165 | NH1 | H-bond | 2.54 |
|  | S43 | O | R165 | NH2 | H-bond | 2.75 |
|  | E46 | OE2 | K245 | NZ | H-bond | 2.88 |
|  | K106 | NZ | E368 | OE2 | Salt bridge | 2.82 |
|  | K106 | NZ | E368 | OE1 | Salt bridge | 3.05 |
|  | R130 | NH1 | D305 | OD1 | Salt bridge | 3.61 |
|  | R130 | NH1 | D305 | OD2 | Salt bridge | 2.77 |
|  | R130 | NH2 | D305 | OD1 | Salt bridge | 3.61 |
|  | R130 | NH2 | D305 | OD2 | Salt bridge | 2.77 |
|  | K133 | NZ | D304 | OD1 | Salt bridge | 3.62 |
|  | K133 | NZ | D304 | OD2 | Salt bridge | 2.40 |
|  | E46 | OE1 | K245 | NZ | Salt bridge | 3.76 |
|  | E46 | OE2 | K245 | NZ | Salt bridge | 3.76 |
| Rpn2 | S469 | OG | D22 | OD1 | H-bond | 2.64 |
|  | N470 | N | D22 | OD1 | H-bond | 3.78 |
|  | N470 | ND2 | D22 | OD1 | H-bond | 3.25 |
|  | R474 | NH2 | D25 | OD2 | H-bond | 3.10 |
|  | N467 | OD1 | N29 | ND2 | H-bond | 3.40 |
|  | D504 | OD1 | R343 | NE | H-bond | 3.13 |
|  | D504 | OD1 | R343 | NH2 | H-bond | 3.07 |
|  | R474 | NH2 | D25 | OD2 | Salt bridge | 3.10 |
|  | D504 | OD2 | R343 | NE | Salt bridge | 3.13 |
|  | D504 | OD2 | R343 | NH1 | Salt bridge | 3.07 |
| Rpt5 | E224 | OE1 | K56 | NZ | H-bond | 2.76 |
|  | E224 | OE2 | S57 | OG | H-bond | 2.81 |
|  | D260 | O | H64 | NE2 | H-bond | 2.75 |
|  | D260 | OD2 | K72 | NZ | H-bond | 2.91 |
|  | E224 | OE2 | K56 | NZ | Salt bridge | 3.70 |
|  | E224 | OE1 | K56 | NZ | Salt bridge | 2.76 |
|  | D260 | OD2 | K72 | NZ | Salt bridge | 2.91 |

